# A plausible identifiable model of the canonical NF-*κ*B signaling pathway

**DOI:** 10.1101/2022.06.09.495460

**Authors:** Joanna Jaruszewicz-Błońska, Ilona Kosiuk, Wiktor Prus, Tomasz Lipniacki

## Abstract

An overwhelming majority of mathematical models of regulatory pathways, including intensively studied NF-*κ*B pathway, remains non-identifiable meaning that their parameters may not be determined by existing data. The existing NF-*κ*B models that are capable to reproduce experimental data, contain non-identifiable parameters, while simplified models with a smaller number of parameters exhibit dynamics that significantly differs from that observed in experiments. Here, we reduce an existing model of the canonical NF-*κ*B pathway by decreasing the number of equations from 15 to 6 in a way that the resulting model exhibits dynamics closely following that of the original model, both for the nominal and the randomly selected parameters. We carried out the sensitivity-based linear analysis and Monte Carlo-based analysis to demonstrate that the resulting model is structurally and practically identifiable based on a simple TNF stimulation protocol in which 5 model variables are measured. The reduced model is capable to reproduce different types of responses characteristic to regulatory motives controlled by negative feedback loops: nearly-perfect adaptation, damped and sustained oscillations. It can serve as a building block of more comprehensive models of immune responses and cancer, where NF-*κ*B plays a decisive role. Our approach, although may not be automatically generalized, suggests that other regulatory pathways models can be transformed to identifiable, while retaining their dynamical features.

## Introduction

The key and frequent drawback of mechanistic models of regulatory pathways is lack of identifiability meaning that model parameters may not be determined by existing data. Here, we focus on the NF-*κ*B regulatory pathway, as recent works point out that the known detailed models for this very important pathway are not identifiable [1], [2], [3]. In this study we reduce the original 15-variable NF-*κ*B model developed by Lipniacki et al. [4] in a way that it preserves the dynamics of key variables (in contrast to ealier simplified models [5], [6], [7]), but makes the model structurally and practically identifiable. Our approach is not automatic, but rather modeler driven and thus differs from the algorithmic methods tested previously on the same model [1], [7], [8] or its later variant [9].

NF-*κ*B is an important transcription factor controlling expression of numerous genes regulating immune responses. It is activated in response to TNF or IL1 cytokines stimulation, as well as viral RNA or bacterial LPS [10]. In resting cells, NF-*κ*B forms inactive cytoplasmic complexes mainly with its primary inhibitor I*κ*B*α*. Signals activating NF-*κ*B converge on the cytoplasmic I*κ*B kinase, IKK. Activated IKK phosphorylates I*κ*B*α*, inducing its ubiquitination and rapid proteasomal degradation. I*κ*B*α* degradation allows for NF-*κ*B translocation to the nucleus, where it upregulates transcription of numerous genes including genes of its two inhibitors I*κ*B*α* and A20 genes [4], [11]. The newly synthesized I*κ*B*α* translocates to the nucleus, binds NF-*κ*B and such formed complex is transported to the cytoplasm, which completes the I*κ*B*α* feedback loop. A second level of NF-*κ*B negative regulation involves A20 that attenuates IKK activity by promoting its transformation into the inactive overphosphorylated form. This way A20 allows for accumulation of a primary NF-*κ*B inhibitor I*κ*B*α* [4], [12].

Modeling efforts, which started two decades ago, resulted in the formulation of detailed biochemical models of the NF-*κ*B signaling pathway [4], [13]. Models, supported by live single cell experiments [14], [15], elucidated the role of I*κ*B*α* and A20 mediated negative feedbacks loops in promoting oscillations observed in several cell lines, including SK-N-AS and 3T3 MEFs. These two feedback loops play also an important role in shaping pulsatile NF-*κ*B responses to pulsatile TNF stimulation [16], [17]; as demonstrated A20 deficient cells exhibit prolonged NF-*κ*B activation in response to a TNF pulse [18]. Individual cells have the ability to respond to the pulsatile TNF stimulation with synchronous NF-*κ*B pulses that result in a high expression of NF-*κ*B regulated genes [15], [19]. Pulsatile stimulation was also shown to transmit more information than the continuous one, with rate of order 1 bit/hour [20].

The main goal of this study is to obtain an accurate, practically identifiable model, and suggest a reasonable experiment allowing for a determination of all parameters with a satisfactory accuracy. We found that a simple protocol with a continuous 2 hour long TNF stimulation followed by a 10 hour long TNF washout phase assures not only structural identifiability of the reduced model, but also allows to determine parameters with accuracy comparable to the more complex pulsatile protocols. We evaluated parameter identifiability using sensitivity analysis techniques [21], [22], [23], [24] finding the parameters that may be determined with the lowest accuracy. We verified practical identifiability performing Monte Carlo simulations [25]. Following the standard Monte Carlo procedure, for each of three different measurement noise levels, we generated 50 different trajectories. Next the model was fitted to such simulated experimental trajectories and 50 parameter estimates were obtained. The results indicate that all parameters can be estimated based on noisy data.

Finally, we showed that the reduced model has ability to produce qualitatively distinct dynamical responses to continuous TNF signal. Depending on the assumed parameters it can exhibit a pulse like response, damped oscillations, or sustained oscillations, i.e., the essential characteristics of the negative feedback system. Importantly, our results suggest that other regulatory pathways models can be transformed to identifiable, while retaining their dynamical features.

## Results

### Model reduction, components and equations of the reduced model

The model of Lipniacki et al. [4], hereafter referred to as the original model, incorporates two negative feedback loops, one mediated by I*κ*B*α* directly binding NF-*κ*B and sequestering it in the cytoplasm, and the other involving the inhibition of IKK (kinase responsible for phoshorylating of I*κ*B*α* and targeting it for proteasomal degradation) by the protein A20 (see Fig 1 A). The dynamics of 15-model variables has been described with mass action kinetics. Starting from the original model we retained the core of the two-feedback network and constructed a reduced model still exhibiting qualitatively similar kinetics. The detailed description of all reduction steps is given in Supplementary Text.

**Fig 1.**
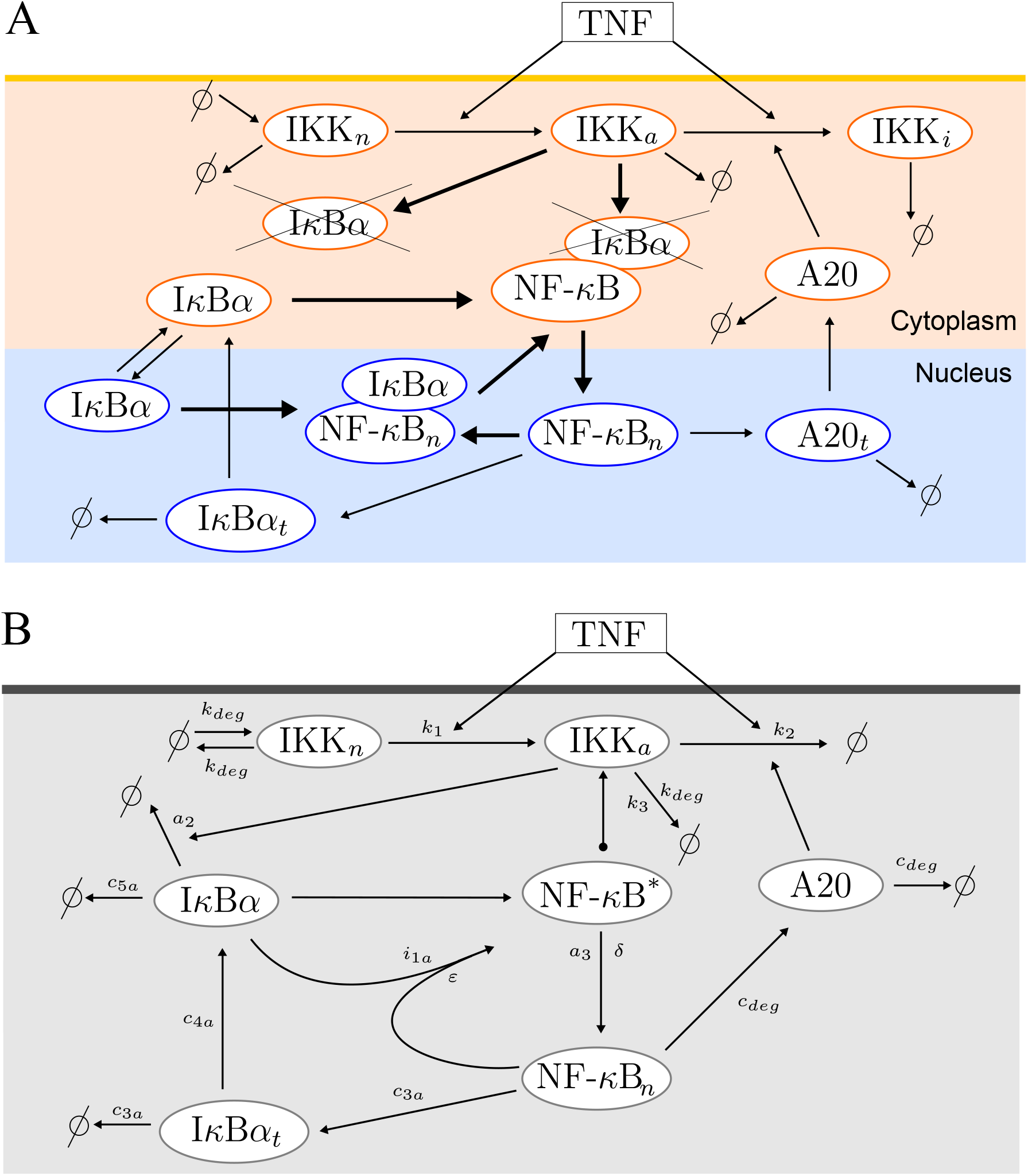
NF-*κ*B pathway diagrams. (A) Diagram of the 15-variable original model. The short lasting cytoplasmic complexes (IKK|I*κ*B*α*) and (IKK|NF-*κ*B|I*κ*B*α*) are not depicted for the sake of simplicity. The bold arrows represent the fast dynamics. (B) Diagram of the reduced 6-variable model. NF-*κ*B* denoting cytoplasmic (NF-*κ*B|I*κ*B*α*) complexes is not associated with the separate equation, since the amount of total NF-*κ*B remains constant: *NFκB_n_* + *NFκB\** = 1.

The key reduction steps can be summarized as follows:

1. We have eliminated the short-lived complexes (IKKa|I*κ*B*α*) and (IKKa|I*κ*B*α*|NF-*κ*B). Instead we equate inflows and outflows rates from these two complexes.
2. We have eliminated cytoplasmic NF-*κ*B (because it either rapidly binds to cytoplasmic I*κ*B*α* or translocates to the nucleus), nuclear I*κ*B*α*(because it rapidly binds to nuclear NF-*κ*B), and the nuclear complexes of (I*κ*B*α*|NF-*κ*B), because they rapidly translocate to the cytoplasm.
3. We have eliminated A20 mRNA, whose formation is a step in A20 protein synthesis. However, we retained I*κ*B*α* mRNA, because we found that the time delay associated with that intermediate step (in contrast to the delay introduced by A20 mRNA) is important for the observed oscillatory behavior.
4. The model was transformed to the nondimensional form. As a consequence of nondimensionalization degradation coefficients for IKKn, A20, and I*κ*B*α* are equal to the corresponding synthesis coefficients (see Table 1).

**Table 1.** Parameter values for the reduced and reduced fitted model.

| Description | Parameter | Pre-fitted value | Fitted value | Unit |
| --- | --- | --- | --- | --- |
| Synthesis and degradation of IKKn | $k_{deg}$ | 0.000125 | 0.000107 | $s^{-1}$ |
| Degradation of IKKa |  |  |  |  |
| Activation of IKKn caused by TNF | $k_1$ | 0.0025 | 0.00195 | $s^{-1}$ |
| Spontaneous inactivation of IKKa | $k_3$ | 0.0015 | 0.00145 | $s^{-1}$ |
| Inactivation of IKKa induced by A20 | $k_2$ | 0.0625 | 0.0357 | $s^{-1}$ |
| IKK ($I\kappa B\alpha$\|NF-κB) association | $a_3$ | 0.2 | 0.0946 | $s^{-1}$ |
| leading to $I\kappa B\alpha$ degradation | | | | |
| Michaelis type constant | $\delta$ | 0.0833 | 0.108 | |
| Michaelis type constant | $\varepsilon$ | 0.0167 | 0.0428 | |
| Synthesis and degradation of A20 | $c_{deg}$ | 0.000171 | 0.000106 | $s^{-1}$ |
| $I\kappa B\alpha$ translation | $c_{4a}$ | 0.0031 | 0.00313 | $s^{-1}$ |
| $I\kappa B\alpha$ degradation induced by IKKa | $a_2$ | 0.04 | 0.0763 | $s^{-1}$ |
| $I\kappa B\alpha$ degradation | $c_{5a}$ | 0.0001 | 0.0000578 | $s^{-1}$ |
| $I\kappa B\alpha$ nuclear import leading to | $i_{1a}$ | 0.001 | 0.000595 | $s^{-1}$ |
| NF-κB removal from the nucleus | | | | |
| $I\kappa B\alpha$ synthesis and degradation of $I\kappa B\alpha$ mRNA | $c_{3a}$ | 0.0004 | 0.000372 | $s^{-1}$ |

The remaining variables are two forms of cytoplasmic IKK (IKKn and IKKa), free nuclear NF-*κ*B, proteins A20 and I*κ*B*α*, and I*κ*B*α* transcript. The resulting model is depicted in Fig 1 B and described by the following system of kinetic equations based on mass action or Michaelis-Menten kinetics.

IKK in its neutral state IKKn. The three terms stand consecutively for IKKn protein synthesis, degradation (with the same coefficient), transition to active IKKa form in response to TNF stimulation (*T_R_* = 1 for TNF ON and *T_R_* = 0 for TNF OFF)

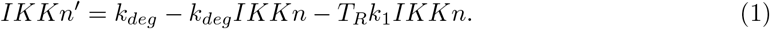

IKK in the active state IKKa. The terms stand consecutively for transition from IKKn form and removal of IKKa (associated with degradation, spontaneous transition to inactive form, and transition to inactive form promoted by A20 and TNF)

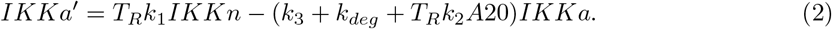

Free nuclear NF-*κ*B. The first term stands for appearance of nuclear NF-*κ*B due to degradation of I*κ*B*α* in (NF-*κ*B|I*κ*B*α*) complexes (whose level is equal to 1 – *NF*κ*B_n_*). The I*κ*B*α* degradation is proportional to IKKa (which phosphorylates I*κ*B*α* and targets it for rapid degradation). The appearance of active NF-*κ*B is blocked by free I*κ*B*α* (that may bind it before its nuclear translocation). The second term describes removal of nuclear NF-*κ*B due to rapid binding with I*κ*B*α*. This process is regulated by I*κ*B*α* nuclear translocation with rate *i_1a_*

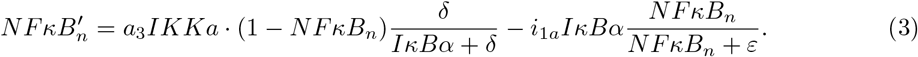

A20 protein. The two terms in the fourth equation describe A20 protein synthesis regulated by NF-*κ*B and its degradation, respectively

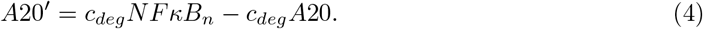

Free cytoplasmic I*κ*B*α* protein. The terms stand consecutively for I*κ*B*α* protein translation, spontaneous degradation, IKKa regulated degradation. The fourth term arises due to reduction of several process: it describes depletion of free I*κ*B*α* due to its binding with free NF-*κ*B that in turn arises due to IKK induced degradation of I*κ*B*α* bounded to NF-*κ*B. The fifth term is equal to the second term in the equation (3) and describes depletion of free I*κ*B*α* due to binding to nuclear NF-*κ*B

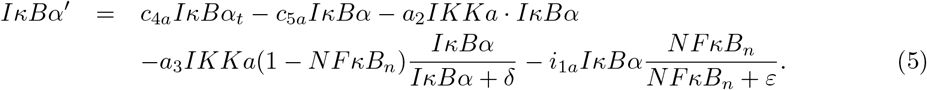

I*κ*B*α* transcript. The terms in the sixth equation describe I*κ*B*α* mRNA synthesis regulated by NF-*κ*B and its degradation, respectively

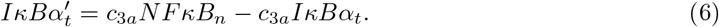

### Refitting the reduced model to the original one

In order to verify that the reduced model may accurately replace the original one, we analyze responses of both models *in silico* experiment involving six TNF stimulation protocols (the tonic stimulation and 5 pulsatile protocols defined in Table S1), which were used in published experiments [16], [17], see Fig 2. In this numerical experiment (hereafter referred to as the combination experiment) we compared numerical trajectories of wild type (WT) and A20 knockouted (A20 KO) cells of the 5 main variables (IKKa, NF-*κ*B, A20, total I*κ*B*α*, and mRNA I*κ*B*α*), generated by the original model, the reduced model and the reduced model fitted to the original one. The combination experiment contains 914 independent data points (defined in Table S1, see also Table S2). The reduced model has been fitted to the original one using PyBioNetFit [26] minimizing squared distances of log-transformed solutions in time points given in Table S1 (see Methods for details). The applied log-transformation follows from the assumption that only the relative (not the absolute) values of all variables are measured. The accuracy of the reduced model and the fitted reduced model with respect to the original one is assessed by the Average Multiplicative Distance (AMD) – see Methods for details. For WT cells the average distance of the reduced model and the original one is AMD_WT_ = 1.55, whereas the distance of the reduced fitted model and the original one is 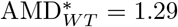 (implying about 30% differences between the two models trajectories). For A20 KO cells these distances are smaller, equal respectively, AMD_A20KO_ = 1.28 and 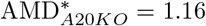. In Fig 2 we show the NF-*κ*B trajectories for the six simulated protocols for the three models; the corresponding trajectories for A20 KO cells are shown in Fig S1. Both figures and the low AMD values between the fitted reduced model and the original one indicate that the reduced model after refitting may accurately replace the original model reproducing trajectories of the 5 main model variables in response to various TNF stimulations.

**Fig 2.**
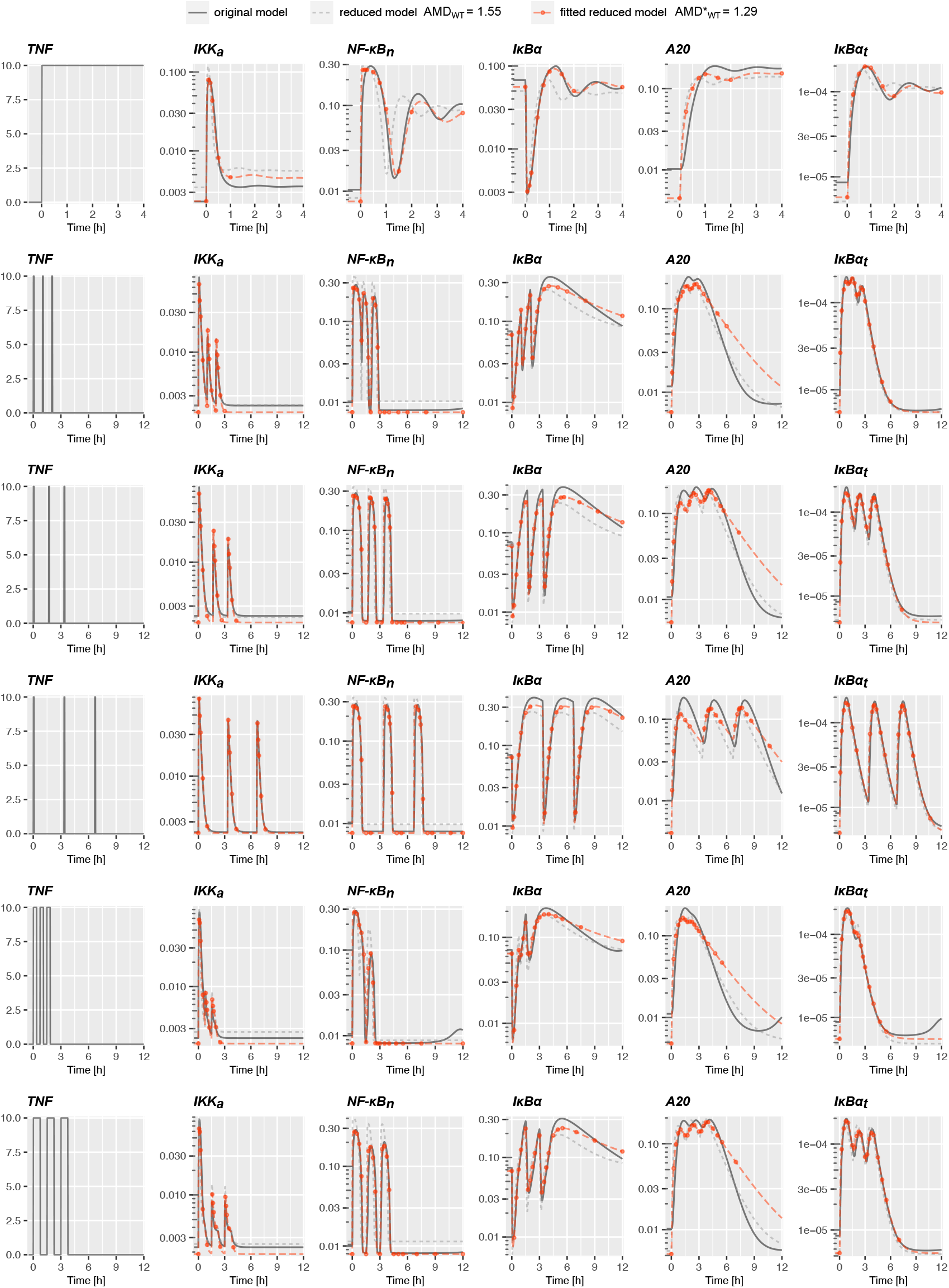
Dynamics of the original, reduced and reduced fitted models in the combination experiment defined in Table S1. Red dots indicate time points from which the nuclear NF-*κ*B and total I*κ*B*α* protein *in silico* measurements are used for fitting the reduced model to the original one. The time points for the remaining variables are given in Table S1.

In Table 1 we provide numerical values of the 13 parameters of the reduced model, before and after refitting them to the original one. The differences between these two sets of parameters are significant, confirming that refitting is an important step after model reduction. With respect to the original model there are two novel parameters: *δ* and *ε*. Parameter *δ* is proportional to NF-*κ*B nuclear import and inversely proportional to I*κ*B*α* – NF-*κ*B binding. Parameter *ε* is proportional to I*κ*B*α* nuclear export and also inversely proportional to I*κ*B*α* and NF-*κ*B binding. Both *δ* and *ε* are smaller than 1, which reflects the original model assumption that the I*κ*B*α*– NF-*κ*B binding is a faster process than nuclear-cytoplasmic trafficking.

The original model has been constrained based on experiments on mouse embryonic fibroblasts. Since the other cell lines may be characterized by different parameters, it is important to verify whether the reduced model can be fitted to the original one also for different sets of parameters. In Fig 3 we have randomly selected five parameter sets of the original model varying the original parameters at most three-fold from their original values (see Table S3) such that the resulting model trajectories vary significantly. Next the reduced model has been fitted to these five new variants of the original model (see Table S4). The comparison of the nuclear NF-*κ*B dynamics of the original and reduced models is shown in Fig 3. For each parameter set the corresponding 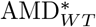 and 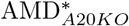 are computed as previously based on the combination experiment. The obtained fits have 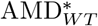 values in range 1.22 - 1.41, and 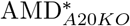 in range 1.13 - 1.33 (The NF-*κ*B trajectories for A20 KO cells for the combination experiment are provided in Fig S2). The small AMD values confirm the ability of the reduced model to reproduce (after the parameter refitting) the original one also for different sets of parameter.

**Fig 3.**
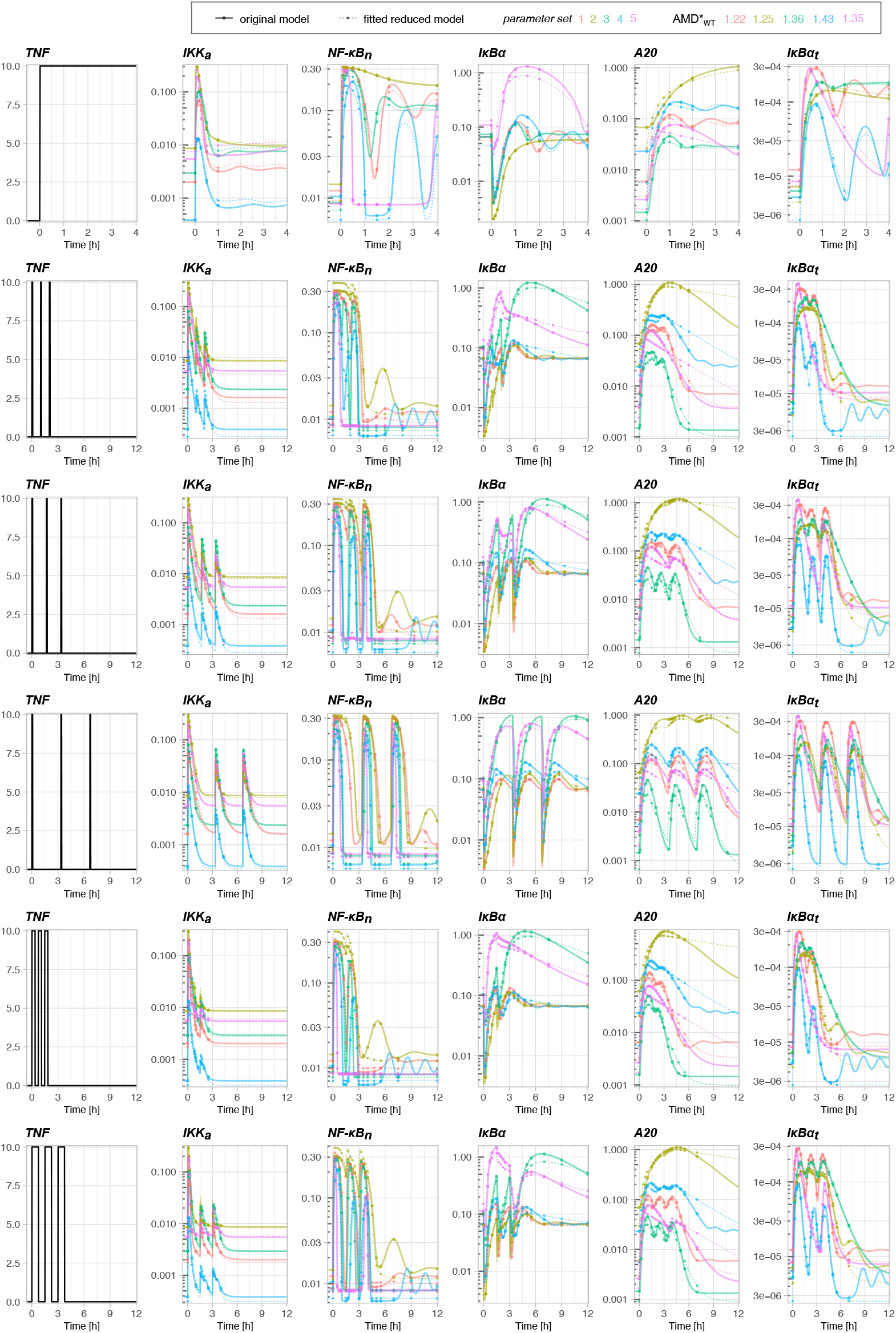
Simulations of the original model for five different sets of parameters and the corresponding reduced model with refitted parameter values (see Table S3 and Table S4). Simulations are performed for the combination experiment defined in Table S1. For each parameter set the corresponding 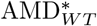 is computed based on trajectories of the 5 main model variables.

In summary, the reduced model may reproduce the NF-*κ*B system dynamics (for WT and A20 KO cells) of the original model both for the original as well as perturbed parameters. Therefore, it represents a good approximation for studying the behavior of the NF-*κ*B signalling pathway. Importantly, we will show that thanks to a smaller number of parameters, the reduced model is, in the contrary to the original one, identifiable.

### Structural identifiability analysis of the original and reduced model

In this section we perform the linear identifiability analysis based on the sensitivity matrix for the original as well as of the reduced model fitted to the original one, hereafter referred as reduced model. In order to demonstrate non-identifiability of the original model based on experiments that have been performed so far [16], [17], we consider the combination experiment (consisting of tonic and 5 pulsatile protocols defined in Table S1). Based on this huge hypothetical experiment (with 914 independent data points), we calculate the sensitivity matrix *S* numerically differentiating log-transformed observables with respect to the log-transformed parameters (as detailed in Methods) for the original model as well as for the reduced model. Based on singular value decomposition of sensitivity matrices we calculate singular values for the original and reduced models. The calculated singular values are scaled by dividing by the squared root of sensitivity vector dimension (*dim*), which assures that they are unchanged by simple repetition of the experiment doubling dim. As shown in Fig 4 for the original model the scaled singular values (hereafter referred as singular values), arranged in descending order, reveal a clear gap indicating the corresponding sensitivity matrix is rank deficient and hence the original model is structurally unidentifiable. In contrast, for the reduced model all singular values are greater than 0.01 indicating structural identifiability.

**Fig 4.**
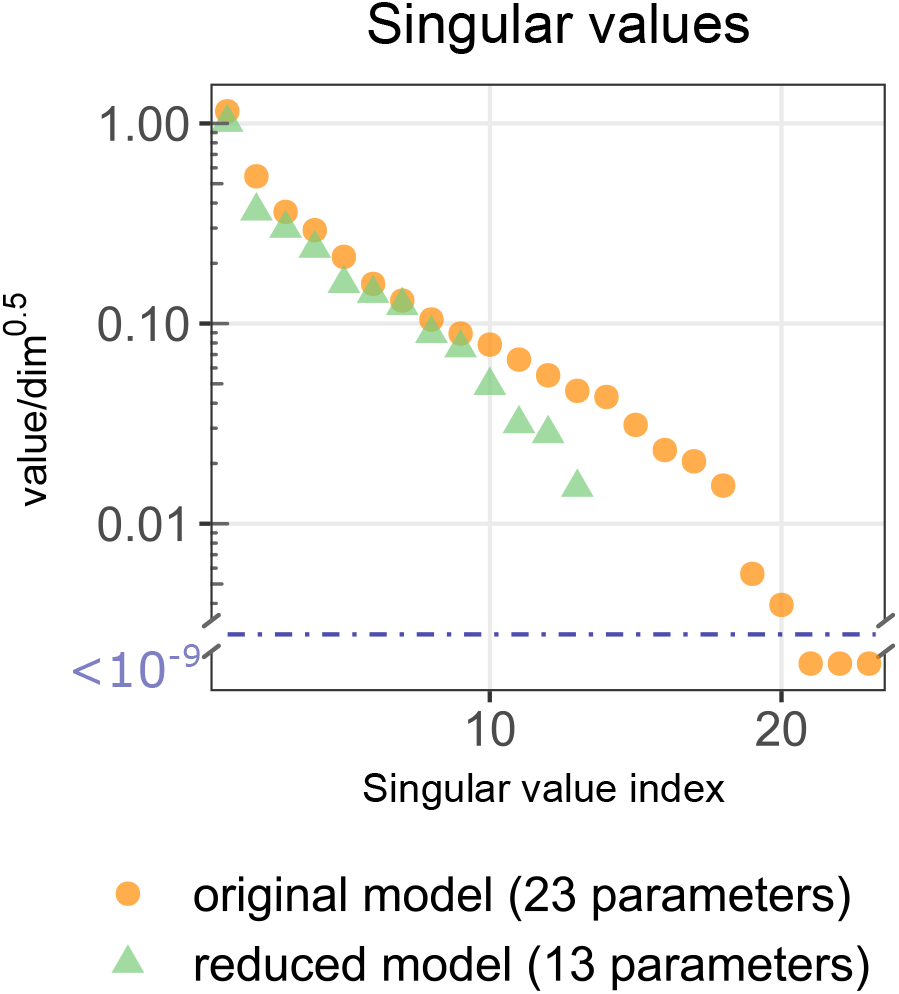
Structural identifiability analysis of the reduced and original models. Singular values of the sensitivity matrices *S* for the original and reduced models. Graphs show all 23 scaled singular values obtained (for the combination experiment) for the original model vs. 13 singular values obtained for the reduced model. Singular values, arranged in descending order, in the original model reveal a clear gap, indicating that the corresponding sensitivity matrix is rank deficient and hence the original model is structurally unidentifiable.

### Detailed linear identifiability analysis of the reduced model

In Fig 5 we evaluate which experimental protocols, defined in Table 2 and Table S1, allow for the most accurate parameter identifiability. In Fig 5 A we show that three smallest singular values for all considered protocols are greater than 10^-3^, indicating that each of protocols renders the reduced model structurally identifiable. We should, however, notice that for the continuous protocol the identifiability is poorest with the smallest singular value equal to 0.0014. For all pulsatile protocols the smallest singular value exceeds 0.007, and when these protocols are combined with the continuous protocol, the smallest singular value rises above 0.014. Interestingly, the simple on-off protocol (proposed in this study, see Table 2) with the smallest number of data points outperforms all remaining protocols having the smallest singular value equal to 0.022. It is important to notice that the comparison of protocols in Fig 5 A is based on scaled singular values, i.e., divided by the squared root of sensitivity vectors dimension. Obviously, the combination experiment having 914 independent data points has greater singular values, than the on-off protocol having only 41 independent data points. The greater scaled singular values of the on-off experiment imply, however, that in order to better identify parameters it pays to repeat the on-off protocol 914/41 ≈ 43 times than do the combination experiment. For the same reason analyzing the identifiability of the model parameters in Figs 5 C and D, and Fig 6 we used the 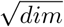 scaled values.

**Fig 5.**
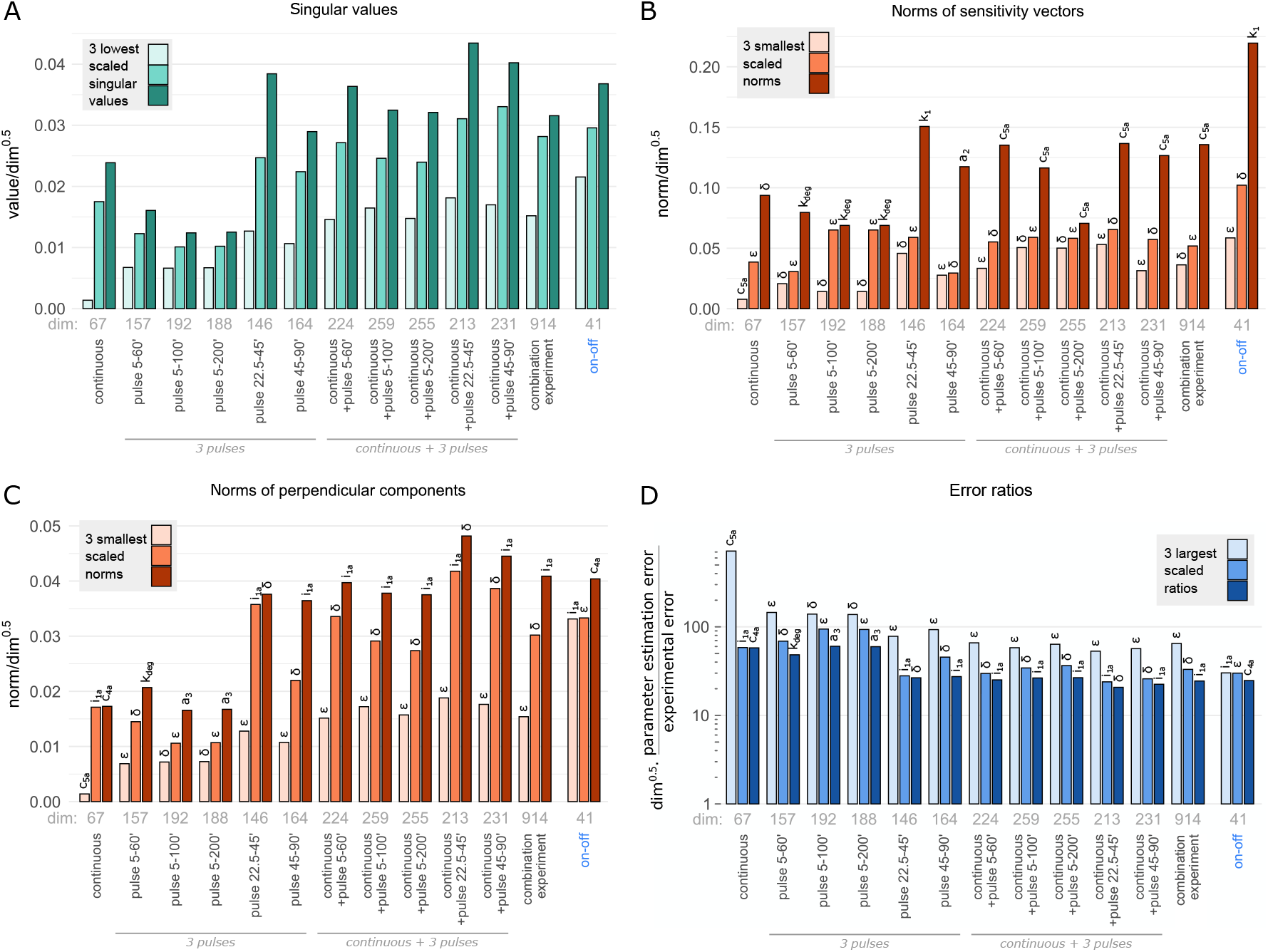
Linear identifiability analysis of the reduced model. (A) Comparison of the three smallest scaled singular values computed for different protocols. (B) Smallest scaled norms of sensitivity vectors ||*S_j_*||. (C) Three smallest scaled norms of perpendicular components 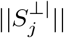. (D) Three largest scaled ratios of the parameter estimation error to experimental error *R_j_* = ln(*σ_linear,j_*)/ln(*σ_data_*). The combination experiment includes the continuous and 5 pulsatile protocols. Scaling with 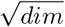 allows to compare the protocols having a different dimension of sensitivity vectors.

**Table 2.** Summary of the on-off protocol of the NF-*κ*B system in WT and A20 KO cells used for *in silico* experiments. A simple protocol with a continuous TNF stimulation (0-120min) followed by a TNF washout (120-720min) for which measured variables are: IKKa, nuclear NF-*κ*B, A20, total I*κ*B*α* protein denoted by I*κ*B*α*\* (*I*κ*B*α*\**=*IκBα*+1 – *NFκB*), and I*κ*B*α* mRNA. For each variable the measurement times used for the sensitivity-based identifiability analysis and reduced model fitting are given.

| on-off | TNF *ON* 0-120 min, TNF *OFF* 120-720 min |
| --- | --- |
| IKKa | 0, 5, 30 min |
| NF-κB | 0, 5, 30, 60, 90, 120, 150, 180 min |
| $I\kappa B\alpha^*$ | 0, 5, 30, 60, 120, 180, 300, 720 min |
| A20, $I\kappa B\alpha_t$ | 0, 30, 60, 300 min |

**Fig 6.**
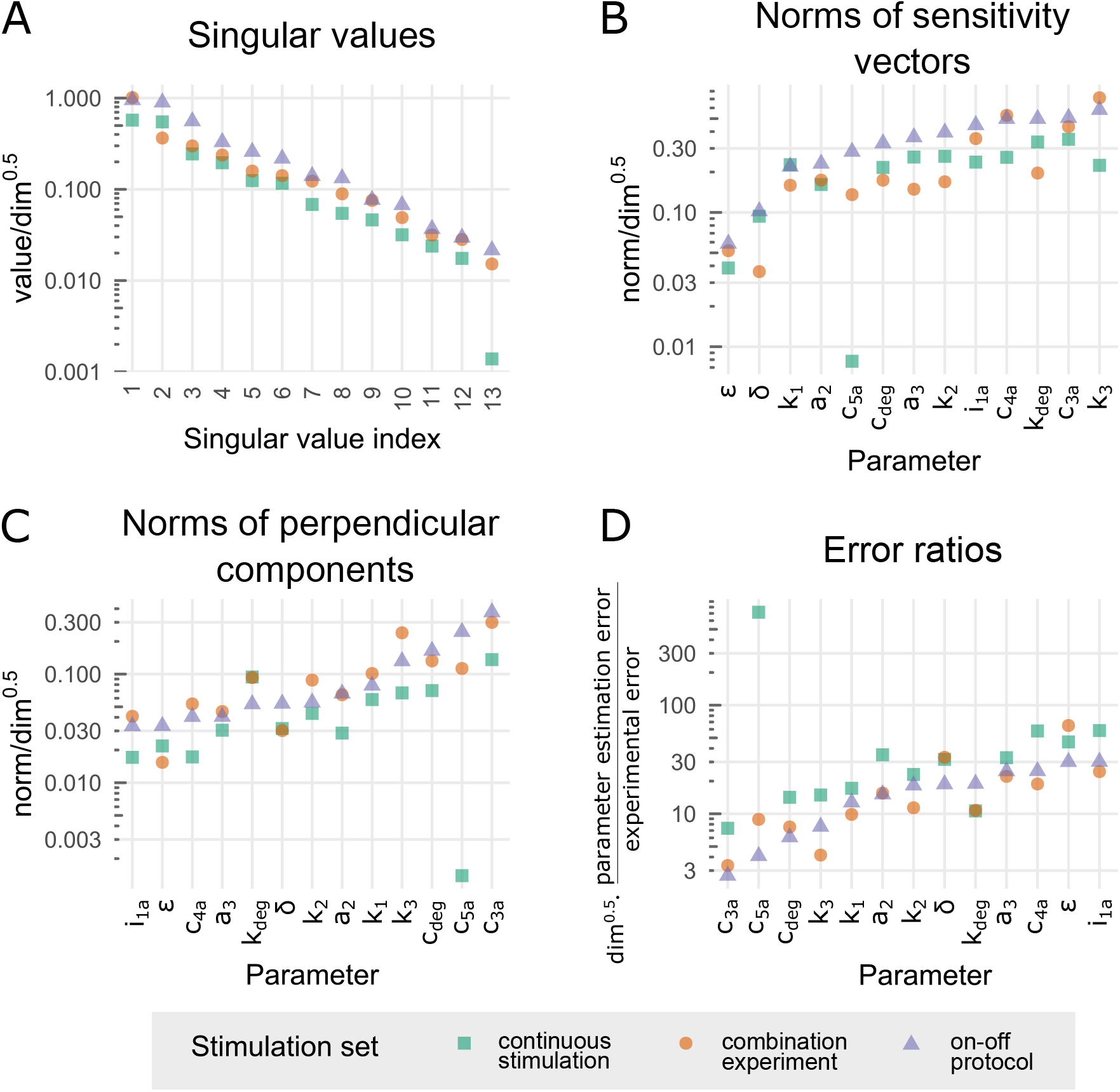
Identifiability analysis of the reduced model - comparison of the protocols: continuous, combination experiment and on-off. (A) Scaled singular values. (B) Scaled norms of sensitivity vectors ||*S_j_*||. (C) Scaled norms of perpendicular components 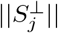. (D) Scaled ratios of parameter estimation error to experimental error ln(*σ_linear,j_*)/ ln(*σ_data_*). Scaling with 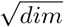 allows for a comparison of the three protocols having a different dimension of sensitivity vectors.

After verifying that the reduced model is structurally identifiable, we further investigate the sensitivity matrix *S* to see which model parameters are the least sensitive and least identifiable for a given protocol. Fig 5 B illustrates the three smallest (scaled) norms corresponding to the least sensitive parameters. It is worth to notice that the parameters *ε* and *δ* appear as two out of three least sensitive parameters for all protocol sets. For the continuous protocol the scaled norm of the sensitivity vector for the parameter *c_5a_* (governing I*κ*B*α* degradation) is the smallest (equal 0.0078) and significantly higher in the on-off protocol (equal to 0.28), which allows to trace I*κ*B*α* degradation.

In Fig 5 C we show the scaled norms of perpendicular parts of sensitivity vectors 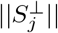 that provide a good measure of identifiability of the associated parameters. A small perpendicular part implies that a change of the associated parameter can be nearly fully compensated by changes of remaining parameters, rendering the considered parameter poorly identifiable. As expected for the continuous protocol the parameter *c_5a_* is least identifiable, however, for the pulsatile protocols the least identifiable are the parameters *δ* and *ε*.

In Fig 5 D we show the three largest scaled ratios of the parameter estimation error to the experimental error. The calculation is based on assumption that all experimental points have lognormally distributed errors with the same geometric standard deviation *σ_data_*. Then the parameter estimation error to the experimental error ratio is equal to ln(*σn_near,j_*)/ln(σ_data_), where *σn_near,j_* is the corresponding geometric standard deviation of parameter estimation error; the subscript ‘linear’ stands for the method of calculation using sensitivity matrix S (see Methods for details). Not surprisingly, for each protocol, the three largest error ratios correspond to the three smallest norms of perpendicular parts 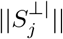 confirming that these parameters are the least identifiable. Figs 5 C and D, show that the on-off protocol out-competes the remaining protocols or their combinations. For this protocol the least identifiable parameter, *i_1a_*, is better identifiable than the least identifiable parameter for any of remaining protocols.

In Fig 6 we focus the identifiability analysis on three particular experiments, i.e., the continuous protocol, the combination experiment, and the on-off protocol. Fig 6 A shows that the simple on-off protocol, out-competes remaining two protocols giving the higher scaled singular values. When the norms of sensitivity vectors Fig 6 B, or their perpendicular components Fig 6 C are juxtaposed, the on-off protocol is comparable to the combination experiment. In Figs 6 B and C, parameters stand in the growing order of respectively norms of sensitivity vectors and their perpendicular components for the on-off protocol. We may notice that the sensitive parameters like *i_1a_* (i.e., having a high norm of sensitivity vector) maybe poorly identifiable, i.e., may have a small perpendicular component (which allows to partially compensate a change of such parameters by respective changes of remaining parameters). A comparison of Figs 6 C and D (with palindromic order of parameters), in turn confirms that parameters with small perpendicular components have a large parameter estimation error to experimental error ratio (and thus may be poorly identifiable based on noisy data).

To sum up, according to our sensitivity-based linear analysis all model parameters are identifiable, however, the continuous protocol results with very poor identifiability of *c_5a_* (governing the I*κ*B*α* degradation). The simple on-off protocol results in comparable or better parameters identifiability than the more complex protocols including the combination experiment.

### Practical identifiability analysis of the reduced model based on Monte Carlo simulations

In this section we verify the practical identifiability of model parameters with help of Monte Carlo simulation [25], [27]. We use the fitted reduced model to simulate the experimental trajectory for the on-off protocol (Fig S3.) Next, we use 50 in *silico* simulated measurements (as defined in Table 2) from the on-off protocol trajectory to refit the model again (using stochastic fitting algorithm; see Methods). Alternatively, we perturb randomly these measurements before refitting. In this latter procedure we mimic the experimental errors [28]. More precisely, we draw values of measurements from the lognormal distribution, Lognormal(*μ*, *σ*^2^), with median *μ* equal to the unperturbed value, and *σ* = ln(*σ_data_*), where *σ_data_* is the assumed geometric standard deviation (sometimes called a multiplicative standard deviation) of measurement values. In Fig 7 we chose three values of *σ_data_*, equal to 1.1, 1.2 and 1.3, and for each *σ_data_* we draw *k* = 50 sets of measurements. For each set of measurements we refit the 13 model parameters. For Fig 7 we select 6 parameters with the highest geometric standard deviation and project k refitted sets of these parameters on the 5 ×6/2 = 15 respective planes. We also show the marginal distribution for each of these 6 parameters. The results for the remaining 7 (better identifiable) parameters are illustrated in Fig S4. Of note 50 *in silico* simulated measurements in the on-off protocol correspond to 41 independent data points, because *n* time points of any time series give n – 1 independent values due to scaling by the geometric average of the series. In the on-off protocol there are 5 series for WT cells for 5 variables, IKKa, NF-*κ*B, A20, total I*κ*B*α*, and mRNA I*κ*B*α*, and 4 series for A20 KO cells for the same variables excluding A20. As a consequence the number of independent data points is by 9 smaller than the number of measurements, see Table S2.

**Fig 7.**
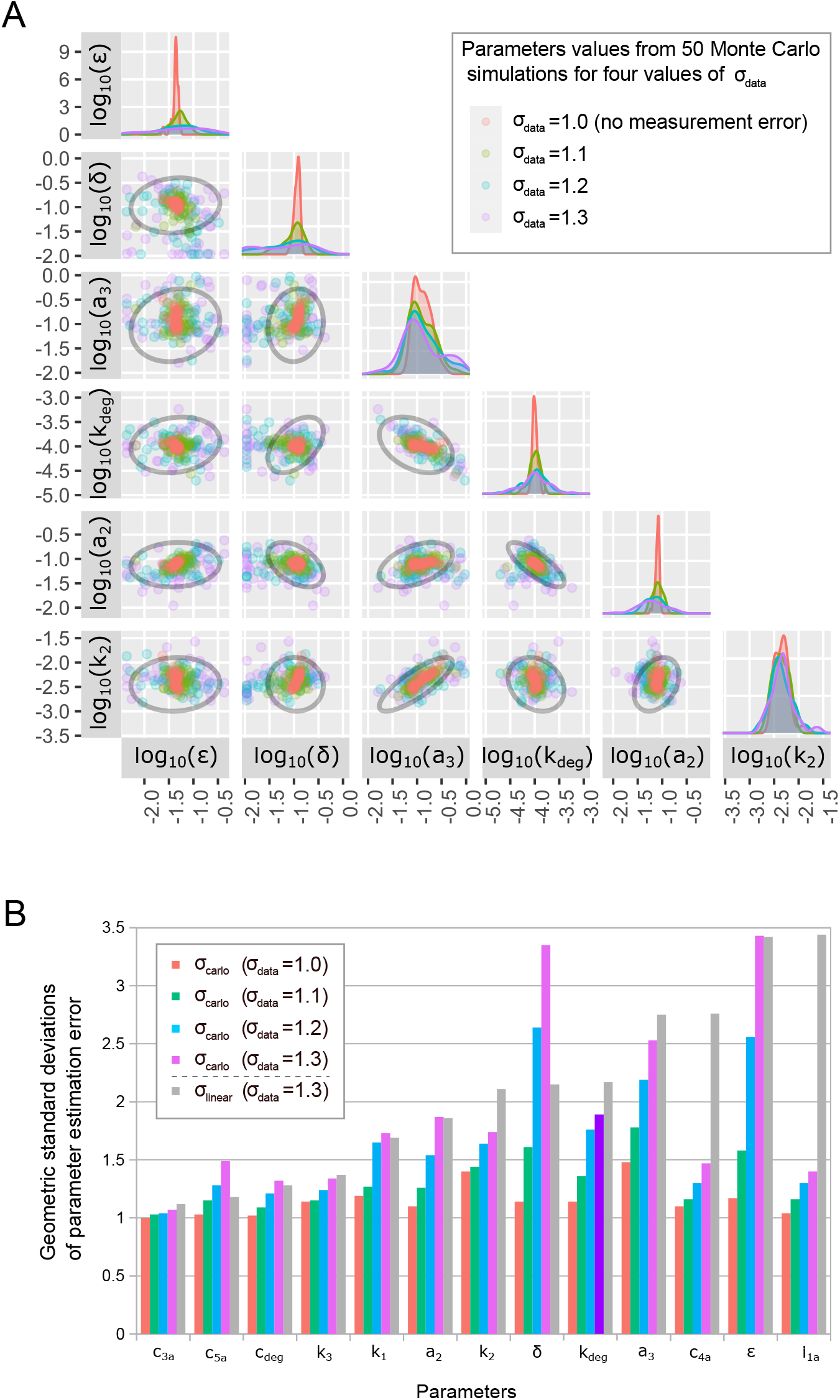
Practical identifiability of the reduced model based on Monte Carlo simulations. (A) A comparison of 75% confidence ellipses (shown in black) computed for *σ_data_* = 1.3 in the linear sensitivity matrix based analysis with results from 50 Monte Carlo simulations for four values of *σ_data_* (1.0, 1.1, 1.2, 1.3). Shown are projections on 15 planes spanned by 6 parameters with the largest geometric standard deviations *σ_carlo,j_* (for *σ_data_* = 1.3). (B) The geometric standard deviation of parameter estimation error *σ_carlo,j_* obtained in Monte Carlo simulation for four values of geometric standard deviations of measurements, *σ_data_* (1.0, 1.1, 1.2, 1.3), is compared with the geometric standard deviation of parameter estimation *σ_linear,j_* obtained from the linear analysis for *σ_data_* = 1.3.

The obtained parameter scatter plots for *σ_data_* = 1.3 are confronted with 75% confidence ellipses obtained from the linear analysis for the same *σ_data_*, showing that parameter uncertainty determined by the linear analysis is comparable to that of determined by Monte Carlo simulations. Fig 7 A shows that due to the use of stochastic fitting algorithm, even when unperturbed measurements are used for *k* independent fittings, the obtained parameters differ from the original ones and correspondingly, their geometric standard deviations *σ_carlo,j_* are greater than 1, see Fig 7 B. As shown in Fig 7 B for all parameters *σ_carlo,j_* grows with *σ_data_*. For most of the parameters the value of *σ_carlo,j_* obtained for *σ_data_* = 1.3 is comparable to the value of *σ_linear,j_* obtained in the linear noise approximation also for *σ_data_* = 1.3, while for some of them (e.g. *δ*) is markedly larger, and a bit surprisingly for some of them (e.g. *i*_1*a*_) is significantly lower. Interestingly, the upper bound of *σ_carlo,j_* and *σ_linear,j_* for all parameters for *σ_data_* = 1.3 is similar, close to 3.5 giving the maximum parameter estimation error to the experimental error ratio equal to ln(3.5)/ln(1.3) ≈4.8 assuring reasonable identifiability of all parameters based on the simple on-off stimulation protocol.

### Characteristic behaviors

In this section we discuss the characteristic dynamical behaviours that can be exhibited by the reduced model (depending on the assumed parameters) in the response to the tonic TNF stimulation. In Fig 8 prior to the tonic TNF stimulation at *t* = 1h, cells remain in the *‘T_R_* = 0’ steady state in which (*IKKn*, *IKKa*, *NFκB_n_*, A20, I*κ*B*α*, I*κ*B*α*_t_) = (1, 0, 0,0,0,0).

**Fig 8.**
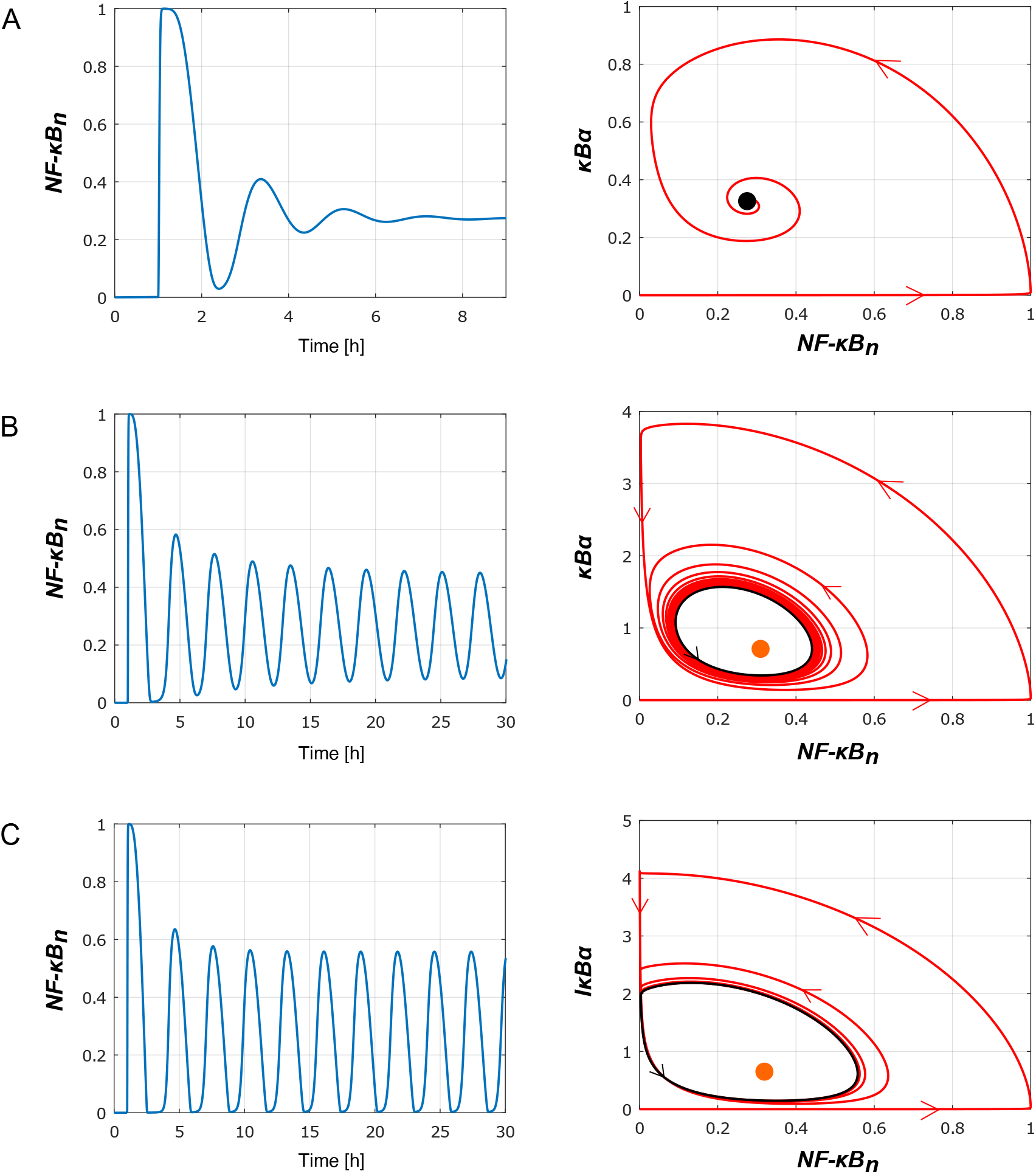
Three types of the oscillatory responses of the reduced model to tonic TFN stimulation: Left subpanels show nuclear *NFκB_n_*(*t*) in response to tonic TNF stimulation started at *t* = 1 hour, whereas the right subpanels show trajectories projected on the (*NFκB_n_*, I*κ*B*α*) plane. (A) Damped oscillations observed for the nominal (fitted to the original model) parameters values (see Table 1), in particular *a*_2_ = 0.0762, *c*_5*a*_ = 0.000058, *i*_1*a*_ = 0.000595 (modified in panels B and C). The black dot on right subpanel shows the ‘*T_R_* = 1’ stable steady state. (B) Limit cycle oscillations of nuclear *NFκB_n_*(*t*) observed in simulations performed for *a_2_* = 0.02, *c_5a_* = 0.00001, *i*_1*a*_ = 0.0001 and other parameters unchanged. The orange dot on the right subpanel shows the ‘*T_R_* = 1’ unstable steady state surrounded by a stable limit cycle (black line). A numerically computed orbit (shown in red) approaches the stable limit cycle. (C) Periodic relaxation-like oscillations observed in simulations performed for *a*_2_ = 0.01, *c*_5*a*_ = 0.00001, *i*_1*a*_ = 0.0001 and other parameters unchanged. Again, the orange dot on the right subpanel shows the ‘*T_R_* = 1’ unstable steady state surrounded by a stable limit cycle (black line).

For the nominal parameter values, i.e., fitted to the original model (see Table 1) the system exhibits dumped oscillations and eventually converges to the *‘T_R_* = 1’ steady state in which all six variables are greater than zero (see Fig 8 A). The dynamics is characterized by a sharp increase of *NF*κ*B_n_* up to 1 (its maximal value). In the short *NF*κ*B_n_* increasing phase, *IκBα* remains close to zero, and then starts increasing after *NFκB_n_* reaches 1, see Fig 8A, right sub-panel.

The observed (dumped) oscillations are a consequence of two negative feedbacks mediated by I*κ*B*α*(directly inhibiting NF-*κ*B) and A20 (inhibiting IKKa that targets I*κ*B*α* for degradation). The I*κ*B*α* feedback is associated with a time delay due to an intermediate step involving I**κ**B*α* transcript, and the process of I*κ*B*α* translocation. Increasing the strength of I*κ*B*α* mediated feedback (by decreasing the I*κ*B*α* degradation coefficients *a*_2_ and *c*_5*a*_) and the associated time delay (by decreasing the I*κ*B*α* nuclear import coefficient *i*_1*a*_), we observe the appearance of limit cycle oscillations, Fig 8 B. As expected, the amplitude of sustained I*κ*B*α* oscillations is much higher (about 3 fold with respect to the nominal parameters, Fig 8 B, right sub-panel), and the period of oscillations is longer (the second *NFκB_n_* peak is at about 5 not 3 hours, Fig 8 B, left sub-panel).

A further decrease of parameter *a*2 causes the growth of the oscillations amplitude, and changes their character, so they resemble spiky oscillations, Fig 8 C. The right subpanel in Fig 8 shows the limit cycle, which consists of one ‘slow’ segment (in which *NFκB_n_* ≈ 0) and one ‘fast’ segment or spike (away from *NFκB_n_* ≈ 0).

Next, we investigate effects of A20 on the NF-*κ*B dynamics. We may observe that after removing A20 negative feedback by setting k_2_=0 the reduced model does not exhibit any oscillations, but after slight overshooting reach ‘*T_R_* = 1’ the steady state characterized by a high level of nuclear *NFκB_n_*, Fig 9 A. This is due to the fact that A20 mediated negative feedback is necessary for functioning of the primary negative feedback of I*κ*B*α*, as only inhibition of IKKa allows for I*κ*B*α* accumulation. Finally, we consider consequences of a dramatic increase of both negative feedbacks resulting from the increase of parameters *k*_2_ and *i*_1*a*_. For the modified parameters we observe a system behavior close to the perfect adaptation [29]. An instant rise of stimulus (from *T*_R_ = 0 to *T*_R_ = 1) results in a spike of *NFκB_n_* followed by a small amplitude damped oscillations converging to the steady state in which *NFκB_n_* is close to zero as in the *‘T_R_* = 0’steady state, Fig 9 B. The perfect adaptation means that the system responds to the temporal change of the stimuli rather than to the actual stimuli amplitude [30].

**Fig 9.**
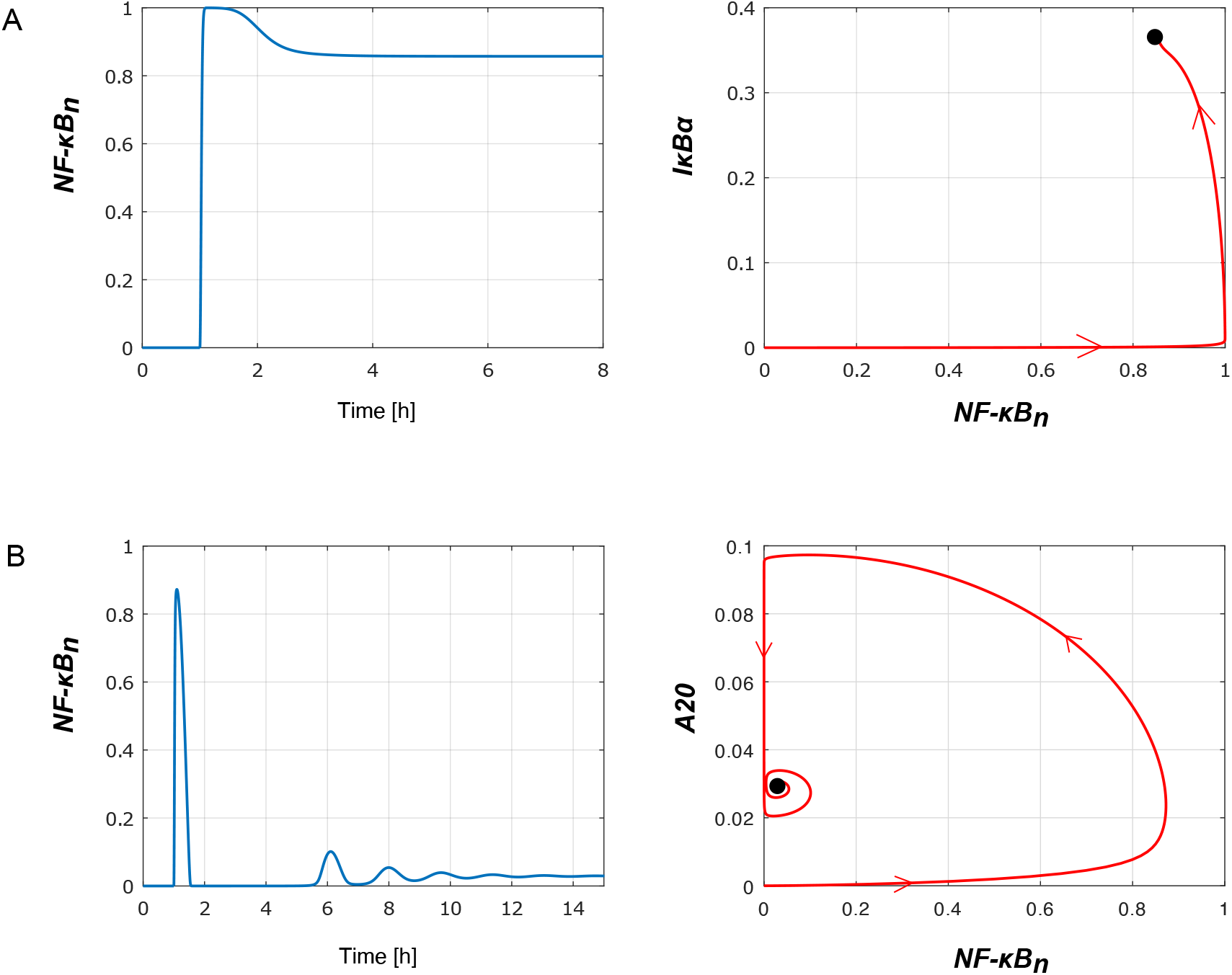
Influence of A20 feedback on the reduced model responses to tonic TNF stimulation. Left subpanels show nuclear *NFκB_n_*(*t*) in response to tonic TNF stimulation started at *t* = 1 hour, whereas the right subpanels show trajectories projected on the (*NFκB_n_,I*κ*B*α**) or (*NFκ*B_n_, A20) planes. (A) Numerically computed solution for *k_2_* =0 (and other parameters unchanged, as in Table 1) implying absence of A20 mediated feedback (as in A20 KO cells). (B) Nearly-perfect adaptation observed in the model variant with significantly stronger negative feedbacks mediated by A20 and *IκBα*. Numerical simulations are performed for *k*_2_ = 3.57 and *i*_1*a*_ = 0.01 (changed from the nominal values *k*_2_ = 0.0357 and *i*_1*a*_ = 0.00595) and other parameters unchanged.

Clearly, *NFκB_n_* dynamics can be dramatically changed by manipulating model parameters. The results presented above indicate that system (1-6) can produce characteristic dynamical responses to a TNF signal: a pulse, damped oscillations, or periodic oscillations (small-amplitude or spike-type). The system is able to respond in a non-adaptive or adaptive way, depending on the strength of negative feedbacks.

## Methods

### Normalization of *in silico* generated data

The applied normalization of the model generated data is based on assumption that experimental measurements are log-normally distributed, i.e., errors are proportional to the measured value of a given variable, provided that this value is above some threshold [28]. Hence, we normalize each simulated time series *x_i_* in two steps

1. replacing each *x_i_* by 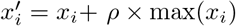, with *ρ* = 0.03.
2. replacing each 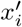 by 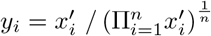.

The first normalization step is needed to avoid values smaller than 0.03 × max (*x_i_*) that typically represent nothing, but measurement noise. The second normalization step reflects the observation that most of the currently available data is in arbitrary units, and thus must be normalized. Normalization by geometric (not arithmetic) mean of time series, reflects the assumption that multiplicative changes of all variables are more important than additive change. An immediate consequence of normalization is that *n* measurements give *n* – 1 independent data points.

### Model fitting

The original and reduced models are both implemented in BioNetGen (cvode solver) and MATLAB. The simulated model outputs are normalized as described above.

In order to fit the reduced model to the original model, we use PyBioNetFit [26], where the corresponding parameter estimation problem in the least-squares sense for the log-transformed systems is formulated [34], [35]. More precisely, the parameter estimation procedure finds a parameter set minimizing the following objective function

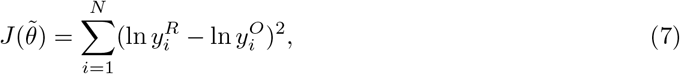

where 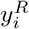 denotes the reduced model outputs, 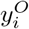 denotes the original model outputs, and *N* is the number of all measurements in the considered protocol. This objective function quantifies the deviation of the (log-transformed) reduced model trajectories from the trajectories of the (log-transformed) original model.

The Average Multiplicative Distance (AMD) between the original and the reduced model (either fitted or not fitted) is defined as

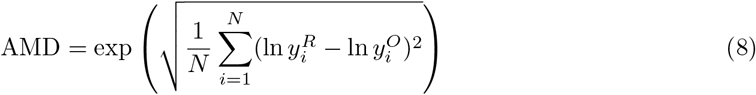

with *N* denoting the number of measurements. We denote by AMD_WT_ and AMD_A20KO_ the average multiplicative distance for WT and A20KO cells, respectively.

### Linear identifiability analysis

For the linear identifiability analysis [36], [37], [38], [39] we use a sensitivity matrix defined as *S* = *S_ij_*, where 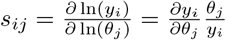 measures a multiplicative change of data value *y_i_* with respect to a multiplicative change of the parameter *θ_j_* (*j* = 1, 2,…, *p* = 13). The matrix *S* is calculated based on the considered stimulation protocols with the corresponding series of measurements of the five reduced model variables (as detailed in Table S1 and Table 2. The elements *s_ij_* of the sensitivity matrix are calculated numerically using the finite differences method [31], with 1% increase/decrease of parameter values. In our analysis, parameters were individually perturbed by a 1% increase/decrease of the original values.

The singular values are calculated by singular value decomposition (SVD) [2], [40]. The perpendicular parts 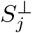 of sensitivity vectors *S_j_* are calculated as 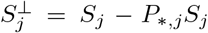. The matrix 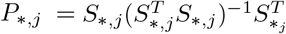 is the so-called projection matrix, where *S*_*,*j*_ denote a matrix formed by removing the *j*-th sensitivity vector from the full sensitivity matrix *S*.

To calculate ratio of parameter estimation error to the experimental error, we assume that experimental data points errors are log-normally distributed, each with the same standard geometric deviation *σ_data_*. As a consequence one may expect that the corresponding parameter estimates are also log-normally distributed with standard geometric deviation *σ_linear,j_*, where *j =1,…,p* is the parameter index. By definition, *R_j_*=ln(*σ_linearj_*)/ln(*σ_data_*). Based on the equation (3.20) in [33], the random variable *ξ_par_* = [*ξ*_*par*,1_,…, *ξ_par,p_*] describing the (linear) parameter estimation error is given by the following formula:

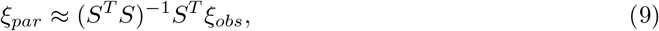

where *ξ_obs_* = [*ξ*_*obs*,1_,…, *ξ_obs,m_*] is a random variable describing the experimental error. Hence, the ratio *R_j_* of the parameter estimation error to the experimental error can be calculated as the *l*_2_-norm of the *j*-th row vector of the *S* pseudo-inverse matrix *A*:=(*S^T^S*)^-1^*S^T^*. We used a mathematically equivalent, but more numerically stable, calculation of *R_j_* based on *QR*-decomposition of *S* (see [34]). In this approach the ratio *R_j_* is given by the *l*_2_-norm of the *i*-th row vector of the 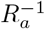 matrix, where *R_a_* denotes the matrix composed of the first *p* rows of an upper triangular matrix *R* from the *QR*-decomposition of the sensitivity matrix *S*.

### Monte Carlo based identifiability analysis

The geometric standard deviation of parameter estimation in Monte Carlo simulations is calculated as

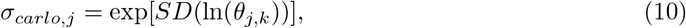

where *SD* denotes a standard deviation, and *θ_j,k_* are *k* = 50 numerical estimates of parameter *θj*,*j* = 1,…, *p* = 13, based on model refitting to the *k* perturbed trajectories.

## Discussion

Mechanistic models of signaling pathways, including the NF-*κ*B pathway, are typically non-identifiable, meaning that their parameters may not be determined based on existing data [41]. In simple words, by fitting a non-identifiable model to experimental data the researcher demonstrates the model ability to reproduce biological observations, but get no or little insight into parameters describing the regulatory process. When a model is structurally non-identifiable, an arbitrary large change of a given parameter can be compensated by an appropriate change of some other parameter (or parameters) rendering this parameter pair (set) non-identifiable. Thus, it is tempting to reduce mechanistic models in such a way that they still perceive dynamics of the observed variables, but reach the stage in which all their parameters become identifiable. Such a reduction involves a number of steps, some being straightforward, while some requiring understanding of model dynamics and can hardly be pursued based on a simple rule based algorithm.

The most obvious step is model nondimensionalization. Indeed, when experimental data give only the relative changes of model variables, the dimensional model may not be identifiable, as it “predicts” absolute values that are not measured. The second step can involve a removal of equations for fast variables assigning the steady state values to corresponding variables - this step requires at least approximated knowledge of time scale of modeled processes. The third step may require a removal of some intermediates that are not experimentally measured. This step is more challenging, as these intermediates contribute to time delays between observed variables. Importantly, a series of intermediates is associated with time delay that follows Erlang type distribution, and thus may not be reproduced by a single process giving exponential distribution of time delay.

In this study we focused on the deterministic model of NF-*κ*B pathway, which contains 15 equations, and in which temporal profiles of nuclear NF-*κ*B are regulated by two interlinked negative feedback loops mediated by I*κ*B*α* and A20. Based on essentially the three outlined above reduction step types, we reduced the considered model to 6 equations parameterized by 13 parameters. We have demonstrated that the reduced model can satisfactorily reproduce the behavior of the original one in response to tonic and five pulsatile experimental protocols (studied before). Moreover, the reduced model can reproduce (after refitting) the original model behavior even when parameters of the original model are randomly changed up to threefold from their original values (which may be needed to reproduce behaviors of different cell types). Next, we demonstrated that the reduced model is structurally and practically identifiable based on experimental protocols and measurements that have been previously applied to the analysis of NF-*κ*B pathway. This implies that the reduced model conserves dynamical properties of the original one, but in contrast to the original model, its parameters can be determined. In particular, we showed, theoretically, that a relatively simple on-off protocol in which 2-hour long TNF stimulation is followed by a 10-hour TNF washed out phase, and in which the main five models variables are measured, suffices to identify model parameters. We verified practical parameter identifiability for the on-off protocol with help of Monte Carlo simulations in which we refitted the model parameters to perturbed observables. The obtained parameter estimation errors follows unimodal distributions with geometric standard deviations comparable to those predicted by the linear analysis based on the sensitivity matrix. This indicates that the reduced model can be fitted to noisy data.

Finally, we demonstrated that the reduced model may exhibit (depending on the assumed parameters regulating negative feedbacks) different types of responses characteristic to regulatory motives controlled by negative feedback loops [42]: nearly-perfect adaptation, damped or sustained oscillations. These responses may allow to reproduce and explain behaviors of different cells types.

In summary, we demonstrated the possibility to reduce the NF-*κ*B pathway model to mdentifiable, but still capable to reproduce rich dynamics of the original model. Our approach is not automatic, as it uses intuitive knowledge about the dynamics of the regulatory system and time scales associated with the involved biochemical processes. Nevertheless, we hope that our study opens a new perspective in modeler-assisted model reduction, and will help to represent other regulatory pathways by identifiable models. Such representation brings the system biology methods closer to those characteristic for physics, where the models, even if approximate, have well defined parameters. The proposed reduced NF-*κ*B pathway model can serve as a building block for more comprehensive models of innate immune responses and cancer, where NF-*κ*B plays a key regulatory role.

## Supporting information

Supplementary Codes - zipped directory

## Data Availability

The computer codes of the computational model developed and analyzed in this study are available as Supplementary Codes.

## Acknowledgments

This research was supported by National Science Centre (Poland) grant 2018/29/B/NZ2/00668 and Norwegian Financial Mechanism GRIEG-1 grant (operated by the National Science Centre, Poland) 2019/34/H/NZ6/00699. Numerical simulations were performed using PLGrid Infrastructure. During the initial stage of the project IK was supported by the European Union’s Horizon 2020 research and innovation program under the Marie Sklodowska-Curie Grant Agreement No 661650. IK thanks Vienna University of Technology for support and hospitality.

## Author Contributions

Conceptualization JJB, IK, WP, TL

Formal Analysis JJB, IK

Methodology JJB, IK

Software JJB

Visualization JJB, IK

Writing – Original Draft JJB, IK, TL

Writing – Review & Editing JJB, IK, WP, TL

## Supporting information

**Supplementary Tables and Figures** Tables S1-S4, Fig. S1-S4

**Supplementary Text** Model reduction, scaling and non-dimensionalization.

**Supplementary Codes** Numerical codes of the computational models.

