## Supplementary Text for "A plausible identifiable model of the canonical NF-*κ*B signaling pathway"

#### Model reduction, scaling and non-dimensionalization

featuring the article

### Model reduction

We will reduce the original model of the NF-$\kappa$B developed by Lipniacki et al.[1] consisting of 15 equations to a 6-variable model. Following the original notation, we use upper-case letters to denote molar concentrations of molecules and their complexes. Nuclear concentrations are denoted by subscript $n$; cytoplasmic concentrations are without any subscript. Concentration of the A20 and I$\kappa$B$\alpha$ transcripts are denoted by subscript $t$.

**Table A. List of the variables of the original model (1).**

| Variable | Definition |
|---|---|
| $IKKn$ | cytoplasmic concentration of neutral form of IKK |
| $IKKa$ | cytoplasmic concentration of active form of IKK |
| $IKKi$ | cytoplasmic concentration of inactive form of IKK |
| $(IKKa\|I\kappa B\alpha)$ | cytoplasmic complex of IKK active and I$\kappa$B$\alpha$ |
| $(IKKa\|I\kappa B\alpha\|NF\kappa B)$ | cytoplasmic complex of IKK active, I$\kappa$B$\alpha$ and NF-$\kappa$B |
| $NF\kappa B$ | cytoplasmic NF-$\kappa$B |
| $NF\kappa B_n$ | nuclear NF-$\kappa$B |
| $A20$ | cytoplasmic A20 protein |
| $A20_t$ | A20 transcript |
| $I\kappa B\alpha$ | cytoplasmic I$\kappa$B$\alpha$ |
| $I\kappa B\alpha_n$ | nuclear I$\kappa$B$\alpha$ |
| $I\kappa B\alpha_t$ | I$\kappa$B$\alpha$ transcript |
| $(I\kappa B\alpha\|NF\kappa B)$ | cytoplasmic complex of I$\kappa$B$\alpha$ and NF-$\kappa$B |
| $(I\kappa B\alpha\|NF\kappa B)_n$ | nuclear complex of I$\kappa$B$\alpha$ and NF-$\kappa$B |
| $Cg$ | control early gene |

The dynamics of the 15 variables listed in Table A is governed by the following system of equations

[1] Lipniacki T, Paszek P, Brasier AR, Luxon B, Kimmel M. Mathematical model of NF-$\kappa$B regulatory module. J. Theor. Biol. 2004; 228(2): 195–215. doi: 10.1016/j.jtbi.2004.01.001. PMID: 15094015.

$$IKKn' = k_{prod} - (k_{deg} + T_R k_1) IKKn, \tag{1.1}$$

$$\begin{aligned} IKKa' = {} & T_R k_1 IKKn - k_3 IKK_a - T_R k_2 IKKa \cdot A20 \\ & - k_{deg} IKKa - a_2 IKKa \cdot I\kappa B\alpha + t_1 (IKKa|I\kappa B\alpha) \\ & - a_3 IKKa \cdot (I\kappa B\alpha|NF\kappa B) + t_2 (IKKa|I\kappa B\alpha|NF\kappa B) \end{aligned} \tag{1.2}$$

$$IKKi' = k_3 IKKa + T_R k_2 IKKa \cdot A20 - k_{deg} IKKi \tag{1.3}$$

$$(IKKa|I\kappa B\alpha)' = a_2 IKKa \cdot I\kappa B\alpha - t_1 (IKKa|I\kappa B\alpha) \tag{1.4}$$

$$\begin{aligned} (IKKa|I\kappa B\alpha|NF\kappa B)' = {} & a_3 IKKa \cdot (I\kappa B\alpha|NF\kappa B) \\ & - t_2 (IKKa|I\kappa B\alpha|NF\kappa B) \end{aligned} \tag{1.5}$$

$$\begin{aligned} NF\kappa B' = {} & c_{6a} (I\kappa B\alpha|NF\kappa B) - a_1 NF\kappa B \cdot I\kappa B\alpha \\ & + t_2 (IKKa|I\kappa B\alpha|NF\kappa B) - i_1 NF\kappa B, \end{aligned} \tag{1.6}$$

$$NF\kappa B_n' = i_1 k_v NF\kappa B - a_1 NF\kappa B_n \cdot I\kappa B\alpha_n \tag{1.7}$$

$$A20' = c_4 A20_t - c_5 A20 \tag{1.8}$$

$$A20_t' = c_2 + c_1 NF\kappa B_n - c_3 A20_t \tag{1.9}$$

$$\begin{aligned} I\kappa B\alpha' = {} & -a_2 IKKa \cdot I\kappa B\alpha - a_1 NF\kappa B \cdot I\kappa B\alpha \\ & + c_{4a} I\kappa B\alpha_t - c_{5a} I\kappa B\alpha - i_{1a} I\kappa B\alpha + e_{1a} I\kappa B\alpha_n \end{aligned} \tag{1.10}$$

$$I\kappa B\alpha_n' = -a_1 NF\kappa B_n \cdot I\kappa B\alpha_n + i_{1a} k_v I\kappa B\alpha - e_{1a} k_v I\kappa B\alpha_n \tag{1.11}$$

$$I\kappa B\alpha_t' = c_{1a} NF\kappa B_n - c_{3a} I\kappa B\alpha_t \tag{1.12}$$

$$\begin{aligned} (I\kappa B\alpha|NF\kappa B)' = {} & a_1 NF\kappa B \cdot I\kappa B\alpha - c_{6a} (I\kappa B\alpha|NF\kappa B) \\ & - a_3 IKKa \cdot (I\kappa B\alpha|NF\kappa B) + e_{2a} (I\kappa B\alpha|NF\kappa B)_n \end{aligned} \tag{1.13}$$

$$(I\kappa B\alpha|NF\kappa B)_n' = a_1 NF\kappa B_n \cdot I\kappa B\alpha_n - e_{2a} k_v (I\kappa B\alpha|NF\kappa B)_n \tag{1.14}$$

$$Cg' = c_{2c} + c_{1c} NF\kappa B_n - c_{3c} Cg. \tag{1.15}$$

The original parameter values are given in Table B.

**Table B. Parameters of the original model (1) and their default numerical values.**

| Parameter | Value | Unit | Description |
|---|---|---|---|
| $C_T$ | 0.3 | $\mu$M | total NF-$\kappa$B concentration w/r/t nuclear volume |
| $k_v$ | 5 | | ratio of cytoplasmic to nuclear volume |
| $c_1$ | 0.0000005 | $s^{-1}$ | inducible A20 mRNA synthesis |
| $c_2$ | 0.0 | $\mu$M$s^{-1}$ | constitutive A20 mRNA synthesis |
| $c_3$ | 0.0004 | $s^{-1}$ | A20 mRNA degradation |
| $c_4$ | 0.5 | $s^{-1}$ | A20 translation |
| $c_5$ | 0.0003 | $s^{-1}$ | A20 degradation |
| $k_1$ | 0.0025 | $s^{-1}$ | IKK activation caused by TNF |
| $k_2$ | 0.1 | $s^{-1}$ | IKK inactivation caused by $A20$ |
| $k_3$ | 0.0015 | $s^{-1}$ | IKK spontaneous inactivation |
| $k_{prod}$ | 0.000025 | $\mu$M$s^{-1}$ | IKKn production rate |
| $k_{deg}$ | 0.000125 | $s^{-1}$ | degradation of IKKa, IKKn and IKKi |
| $a_2$ | 0.2 | $\mu$M$^{-1}s^{-1}$ | IKK I$\kappa$B$\alpha$ association |
| $a_1$ | 0.5 | $\mu$M$^{-1}s^{-1}$ | I$\kappa$B$\alpha$ NF-$\kappa$B association |
| $a_3$ | 1 | $\mu$M$^{-1}s^{-1}$ | IKK (I$\kappa$B$\alpha$ \| NF-$\kappa$B) association |
| $t_1$ | 0.1 | $s^{-1}$ | degradation of I$\kappa$B$\alpha$ in (IKK \| I$\kappa$B$\alpha$) complex |
| $t_2$ | 0.1 | $s^{-1}$ | degradation of I$\kappa$B$\alpha$ in (IKK \| I$\kappa$B$\alpha$ \| NF-$\kappa$B) complex |
| $c_{1a}$ | 0.0000005 | $s^{-1}$ | inducible I$\kappa$B$\alpha$ mRNA synthesis |
| $c_{2a}$ | 0.0 | $\mu$M$s^{-1}$ | constitutive mRNA I$\kappa$B$\alpha$ synthesis |
| $c_{3a}$ | 0.0004 | $s^{-1}$ | I$\kappa$B$\alpha$ mRNA degradation |
| $c_{4a}$ | 0.5 | $s^{-1}$ | I$\kappa$B$\alpha$ translation |
| $c_{5a}$ | 0.0001 | $s^{-1}$ | I$\kappa$B$\alpha$ degradation |
| $c_{6a}$ | 0.00002 | $s^{-1}$ | degradation I$\kappa$B$\alpha$ in (I$\kappa$B$\alpha$ \| NF-$\kappa$B) complex |
| $i_1$ | 0.0025 | $s^{-1}$ | NF-$\kappa$B nuclear import |
| $e_{2a}$ | 0.01 | $s^{-1}$ | (I$\kappa$B$\alpha$ \| NF-$\kappa$B) nuclear export |
| $i_{1a}$ | 0.001 | $s^{-1}$ | I$\kappa$B$\alpha$ nuclear import |
| $e_{1a}$ | 0.0005 | $s^{-1}$ | I$\kappa$B$\alpha$ nuclear export |
| $c_{1c}$ | 0.0000005 | $s^{-1}$ | inducible transcription |
| $c_{2c}$ | 0. | $\mu$M$s^{-1}$ | constitutive transcription |
| $c_{3c}$ | 0.0004 | $s^{-1}$ | control gene mRNA degradation |

The model reduction is carried out in six steps discussed below.

**Step 1**. Elimination of equations for $Cg$ and $IKKi$. Elimination of terms with zero rates.

We eliminate $Cg$ and $IKKi$ as non-essential readouts not influencing the dynamics of the remaining 13 variables. We eliminate terms having in the original model (1) zero rates, i.e., $c_2, c_{2a}, c_{2c}$. In addition, we set $c_{6a} = 0$.

**Step 2**. Elimination of equations (1.4) and (1.5) describing the kinetics of $(IKKa|I\kappa B\alpha)$ and $(IKKa|I\kappa B\alpha|NF\kappa B)$ complexes.

In this step, as well as in steps 3 and 4, we eliminate fast variables, which describe either short lasting complexes or proteins that either rapidly bind to other proteins or are rapidly translocated to the other compartment.

Binding of IKKa to $I\kappa B\alpha$ and $(I\kappa B\alpha|NF\kappa B)$ results in $I\kappa B\alpha$ degradation. Since $t_1 = t_2 = 0.1\ s^{-1}$ complexes $(IKKa|I\kappa B\alpha)$ and $(IKKa|I\kappa B\alpha|NF\kappa B)$ are short lasting, with repect to the characteristic scale of regulatory process. Thus, one can neglect these intermediate steps in $I\kappa B\alpha$ degradation and assume that IKKa catalyzes $I\kappa B\alpha$ degradation with rates $a_2$ and $a_3$ equal to the rates of complex formation, which leads to the following 11-dimensional system

$$IKKn' = k_{prod} - (k_{deg} + T_R k_1) IKKn, \tag{2.1}$$

$$IKKa' = T_R k_1 IKKn - k_3 IKKa - T_R k_2 IKKa \cdot A20 - k_{deg} IKKa, \tag{2.2}$$

$$NF\kappa B' = a_3 IKKa \cdot (I\kappa B\alpha|NF\kappa B) - a_1 NF\kappa B \cdot I\kappa B\alpha - i_1 NF\kappa B, \tag{2.3}$$

$$NF\kappa B_n' = i_1 k_v NF\kappa B - a_1 NF\kappa B_n \cdot I\kappa B\alpha_n, \tag{2.4}$$

$$A20' = c_4 A20_t - c_5 A20, \tag{2.5}$$

$$A20_t' = c_1 NF\kappa B_n - c_3 A20_t, \tag{2.6}$$

$$\begin{aligned} I\kappa B\alpha' = & -a_2 IKKa \cdot I\kappa B\alpha - a_1 NF\kappa B \cdot I\kappa B\alpha + c_{4a} I\kappa B\alpha_t \\ & - c_{5a} I\kappa B\alpha - i_{1a} I\kappa B\alpha + e_{1a} I\kappa B\alpha_n, \end{aligned} \tag{2.7}$$

$$I\kappa B\alpha_n' = i_{1a} k_v I\kappa B\alpha - a_1 NF\kappa B_n \cdot I\kappa B\alpha_n - e_{1a} k_v I\kappa B\alpha_n, \tag{2.8}$$

$$I\kappa B\alpha_t' = c_{1a} NF\kappa B_n - c_{3a} I\kappa B\alpha_t, \tag{2.9}$$

$$\begin{aligned} (I\kappa B\alpha|NF\kappa B)' = & \ a_1 NF\kappa B \cdot I\kappa B\alpha - a_3 IKKa \cdot (I\kappa B\alpha|NF\kappa B) \\ & + e_{2a} (I\kappa B\alpha|NF\kappa B)_n, \end{aligned} \tag{2.10}$$

$$(I\kappa B\alpha|NF\kappa B)_n' = a_1 NF\kappa B_n \cdot I\kappa B\alpha_n - e_{2a} k_v (I\kappa B\alpha|NF\kappa B)_n. \tag{2.11}$$

**Step 3**. Elimination of the equations for cytoplasmic $NF\kappa B$ and nuclear $I\kappa B\alpha_n$

In the equation (2.3) for the cytoplasmic $NF\kappa B$ the first term represents liberation NF-$\kappa$B due to degradation of I$\kappa$B$\alpha$ in the cytoplasmic $(I\kappa B\alpha|NF\kappa B)$ complexes, the second term represents $NF\kappa B$ depletion due to formation of complexes with I$\kappa$B$\alpha$. The last term describes transport of free cytoplasmic NF-$\kappa$B to the nucleus. As these two last processes are fast we may in approximation neglect $NF\kappa B'$. We assume the steady state of free cytoplasmic NF-$\kappa$B is reached ($NF\kappa B' = 0$) and thus obtain

$$NF\kappa B = \frac{a_3 IKKa \cdot (I\kappa B\alpha|NF\kappa B)}{a_1 I\kappa B\alpha + i_1}. \tag{3}$$

In the equation (2.8) for the nuclear free $I\kappa B\alpha_n$ the first term describes I$\kappa$B$\alpha$ transport to the nucleus, the second term describes depletion of $I\kappa B\alpha_n$ due to formation of the nuclear complexes $(I\kappa B\alpha|NF\kappa B)_n$, while the last term represent the I$\kappa$B$\alpha$ transport out of the nucleus. Similarly, because these two last processes are fast, we will in approximation neglect $I\kappa B\alpha_n'$ and thus obtain

$$I\kappa B\alpha_n = \frac{i_{1a}k_v I\kappa B\alpha}{a_1 NF\kappa B_n + e_{1a}k_v}. \tag{4}$$

Consequently, using (3) and (4), we obtain the following system of 9 equations

$$IKKn' = k_{prod} - (k_{deg} + T_R k_1) IKKn, \tag{5.1}$$

$$IKKa' = T_R k_1 IKKn - k_3 IKKa - T_R k_2 IKKa \cdot A20 - k_{deg} IKKa, \tag{5.2}$$

$$NF\kappa B_n' = k_v a_3 IKKa \cdot (I\kappa B\alpha|NF\kappa B)\frac{\mu}{I\kappa B\alpha + \mu} - i_{1a}k_v I\kappa B\alpha \frac{NF\kappa B_n}{NF\kappa B_n + \nu}, \tag{5.3}$$

$$A20' = c_4 A20_t - c_5 A20, \tag{5.4}$$

$$A20_t' = c_1 NF\kappa B_n - c_3 A20_t, \tag{5.5}$$

$$\begin{aligned} I\kappa B\alpha' = {} & c_{4a} I\kappa B\alpha_t - a_2 IKKa \cdot I\kappa B\alpha - c_{5a} I\kappa B\alpha \\ & - a_3 IKKa \cdot (I\kappa B\alpha|NF\kappa B)\frac{I\kappa B\alpha}{I\kappa B\alpha + \mu} - i_{1a} I\kappa B\alpha \frac{NF\kappa B_n}{NF\kappa B_n + \nu}, \end{aligned} \tag{5.6}$$

$$I\kappa B\alpha_t' = c_{1a} NF\kappa B_n - c_{3a} I\kappa B\alpha_t, \tag{5.7}$$

$$(I\kappa B\alpha|NF\kappa B)' = - a_3 IKKa \cdot (I\kappa B\alpha|NF\kappa B)\frac{\mu}{I\kappa B\alpha + \mu} + e_{2a}(I\kappa B\alpha|NF\kappa B)_n, \tag{5.8}$$

$$(I\kappa B\alpha|NF\kappa B)_n' = i_{1a}k_v I\kappa B\alpha \frac{NF\kappa B_n}{NF\kappa B_n + \nu} - e_{2a}k_v (I\kappa B\alpha|NF\kappa B)_n, \tag{5.9}$$

where $\mu := \frac{i_1}{a_1}$ and $\nu := \frac{e_{1a}k_v}{a_1}$.

**Step 4**. Elimination of equation (5.9) for the nuclear complex $(I\kappa B\alpha|NF\kappa B)_n$

The equation (5.9) contains two terms: the first one describes formation of $(I\kappa B\alpha|NF\kappa B)_n$ nuclear complexes and the second one describes their transport out of the nucleus. Since these processes are relatively fast in approximation we neglect $(I\kappa B\alpha|NF\kappa B)_n'$, and thus obtain $e_{2a}k_v(I\kappa B\alpha|NF\kappa B)_n = i_{1a}k_v I\kappa B\alpha \frac{NF\kappa B_n}{NF\kappa B_n + \nu}$. This allows us to eliminate equation (5.9) and replace equation (5.8) by

$$(I\kappa B\alpha|NF\kappa B)' = -a_3 IKKa \cdot (I\kappa B\alpha|NF\kappa B)\frac{\mu}{I\kappa B\alpha + \mu} + i_{1a} I\kappa B\alpha \frac{NF\kappa B_n}{NF\kappa B_n + \nu}. \tag{6}$$

**Step 5**. Elimination of cytoplasmic complex $(I\kappa B\alpha|NF\kappa B)$.

The resulting 8-variable system (5.1)–(5.7) and (6) has one conservation law: $NF\kappa B_n' + k_v(I\kappa B\alpha|NF\kappa B)' = 0$. Hence, we obtain $(I\kappa B\alpha|NF\kappa B) = (C_T - NF\kappa B_n)/k_v$ with the maximal concentration of nuclear NF-$\kappa$B denoted by $C_T$=0.3$\mu$M. This in turn corresponds to the cytoplasmic concentration of 0.06$\mu$M, as it is assumed that the cytoplasmic volume is $k_v = 5$ times larger. Note that two proteins I$\kappa$B$\alpha$ and NF-$\kappa$B may form complexes in the cytoplasm, and although the variable describing NF-$\kappa$B | I$\kappa$B$\alpha$ complexes will not explicitly be present in the reduced model, the concentration of these complexes is given by $(C_T - NF\kappa B_n)/k_v$.

The resulting reduced 7-variable system becomes

$$IKKn' = k_{prod} - (k_{deg} + T_R k_1) IKKn, \tag{7.1}$$

$$IKKa' = T_R k_1 IKKn - k_3 IKKa - T_R k_2 IKKa \cdot A20 - k_{deg} IKKa, \tag{7.2}$$

$$NF\kappa B_n' = a_3 IKKa \cdot (C_T - NF\kappa B_n) \frac{\mu}{I\kappa B\alpha + \mu} - i_{1a} k_v I\kappa B\alpha \frac{NF\kappa B_n}{NF\kappa B_n + \nu}, \tag{7.3}$$

$$A20' = c_4 A20_t - c_5 A20, \tag{7.4}$$

$$A20_t' = c_1 NF\kappa B_n - c_3 A20_t, \tag{7.5}$$

$$\begin{aligned} I\kappa B\alpha' = {} & c_{4a} I\kappa B\alpha_t - a_2 IKKa \cdot I\kappa B\alpha - c_{5a} I\kappa B\alpha \\ & - \frac{a_3}{k_v} IKKa (C_T - NF\kappa B_n) \frac{I\kappa B\alpha}{I\kappa B\alpha + \mu} - i_{1a} I\kappa B\alpha \frac{NF\kappa B_n}{NF\kappa B_n + \nu}, \end{aligned} \tag{7.6}$$

$$I\kappa B\alpha_t' = c_{1a} NF\kappa B_n - c_{3a} I\kappa B\alpha_t. \tag{7.7}$$

**Step 6**. Elimination of $A20_t$

In approximation we replace the equations (7.4)–(7.5) with a single new equation for protein A20

$$A20' = c_{prod} NF\kappa B_n - c_{deg} A20, \tag{8}$$

where $c_{prod} := \frac{c_1 c_4}{(c_3 + c_5)}$ and $c_{deg} := \frac{c_3 c_5}{(c_3 + c_5)}$ are effective coefficients A20 protein synthesis and A20 protein coefficient, respectively. These coefficients assure that for constant $NF\kappa B_n$ the equilibrium A20 protein concentration is

$$A20_{eq} = \frac{c_{prod} NF\kappa B_n}{c_{deg}} = \frac{c_1 c_4 NF\kappa B_n}{c_3 c_5} \tag{9}$$

equal to the equilibrium concentration following from the equations (7.4)–(7.5). Additionally, according to the equations (7.4)–(7.5) the characteristic time of reaching equilibrium concentration for A20 mRNA is $1/c_3$, while the characteristic time in which A20 protein reaches equilibrium concentration is $1/c_5$. A sum of these times is $\frac{c_3 + c_5}{c_3 c_5} = 1/c_{deg}$.

The resulting final reduced system is 6-dimensional and its dynamics is governed by

$$\begin{aligned}
IKKn' &= k_{prod} - (k_{deg} + T_R k_1) IKKn, \\
IKKa' &= T_R k_1 IKKn - k_3 IKKa - T_R k_2 IKKa \cdot A20 - k_{deg} IKKa, \\
NF\kappa B_n' &= a_3 IKKa \cdot (C_T - NF\kappa B_n) \frac{\mu}{I\kappa B\alpha + \mu} - i_{1a} k_v I\kappa B\alpha \frac{NF\kappa B_n}{NF\kappa B_n + \nu}, \\
A20' &= c_{prod} NF\kappa B_n - c_{deg} A20, \qquad (10) \\
I\kappa B\alpha' &= c_{4a} I\kappa B\alpha_t - a_2 IKKa \cdot I\kappa B\alpha - \frac{a_3}{k_v} IKKa (C_T - NF\kappa B_n) \frac{I\kappa B\alpha}{I\kappa B\alpha + \mu} \\
&\quad - c_{5a} I\kappa B\alpha - i_{1a} I\kappa B\alpha \frac{NF\kappa B_n}{NF\kappa B_n + \nu} \\
I\kappa B\alpha_t' &= c_{1a} NF\kappa B_n - c_{3a} I\kappa B\alpha_t.
\end{aligned}$$

The parameters of the reduced model (10) and their numerical values obtained from the model reduction are given in Table D.

**Table C. Variables of the reduced model (10).**

| Variable | Definition |
|---|---|
| $IKKn$ | cytoplasmic concentration of neutral form of IKK |
| $IKKa$ | cytoplasmic concentration of active form of IKK |
| $NF\kappa B_n$ | free nuclear NF-$\kappa$B |
| $A20$ | A20 protein |
| $I\kappa B\alpha$ | free cytoplasmic I$\kappa$B$\alpha$ protein |
| $I\kappa B\alpha_t$ | I$\kappa$B$\alpha$ transcript |

**Table D. Parameters of the reduced model (10) and their numerical values obtained from the model reduction.**

| Parameter | Value | Unit | Description |
|---|---|---|---|
| $C_T$ | 0.3 | $\mu M$ | total NF-$\kappa$B concentration w/r/t nuclear volume |
| $k_v$ | 5 | | ratio of cytoplasmic to nuclear volume |
| $k_1$ | 0.0025 | $s^{-1}$ | IKK activation caused by TNF |
| $k_2$ | 0.1 | $s^{-1}$ | IKK inactivation caused by A20 |
| $k_3$ | 0.0015 | $s^{-1}$ | IKK spontanouous inactivation |
| $k_{prod}$ | 0.000025 | $\mu$M$s^{-1}$ | IKKn production rate |
| $k_{deg}$ | 0.000125 | $s^{-1}$ | degradation of IKKa and IKKn |
| $\mu$ | 0.005 | $\mu$M | Michaelis constant for inhibition of nuclear NF-$\kappa$B by I$\kappa$B$\alpha$ |
| $\nu$ | 0.005 | $\mu$M | Michaelis constant for the export of NF-$\kappa$B from the nucleus |
| $c_{prod}$ | 0.000357 | $s^{-1}$ | effective A20 protein synthesis coefficient |
| $c_{deg}$ | 0.000171 | $s^{-1}$ | effective A20 degradation coefficient |
| $a_2$ | 0.2 | $\mu$M$^{-1}s^{-1}$ | IKK I$\kappa$B$\alpha$ association leading to I$\kappa$B$\alpha$ degradation |
| $a_3$ | 1 | $\mu$M$^{-1}s^{-1}$ | IKK (I$\kappa$B$\alpha$ \| NF-$\kappa$B) association leading to I$\kappa$B$\alpha$ degradation |
| $c_{1a}$ | 0.0000005 | $s^{-1}$ | inducible I$\kappa$B$\alpha$ mRNA synthesis |
| $c_{3a}$ | 0.0004 | $s^{-1}$ | I$\kappa$B$\alpha$ mRNA degradation |
| $c_{4a}$ | 0.5 | $s^{-1}$ | I$\kappa$B$\alpha$ translation |
| $c_{5a}$ | 0.0001 | $s^{-1}$ | I$\kappa$B$\alpha$ degradation |
| $i_{1a}$ | 0.001 | $s^{-1}$ | I$\kappa$B$\alpha$ nuclear import |

### Model scaling and non-dimensionalization

The available data in most cases lacks the absolute quantification. Therefore, the dependent variables must be non-dimensionalized in order to make the model identifiable. We introduce the following scaling leaving time the only dimensional variable.

As a first step, we scale

$$NF\kappa B_n := k_v N\tilde{F}\kappa B, \tag{11}$$

which results in

$$
\begin{aligned}
IKKn' &= k_{prod} - (k_{deg} + T_R k_1) IKKn, \\
IKKa' &= T_R k_1 IKKn - k_3 IKKa - T_R k_2 IKKa \cdot A20 - k_{deg} IKKa, \\
k_v N\tilde{F}\kappa B_n' &= a_3 IKKa \cdot (C_T - k_v N\tilde{F}\kappa B_n) \frac{\mu}{I\kappa B\alpha + \mu} - i_{1a} k_v I\kappa B\alpha \frac{k_v N\tilde{F}\kappa B_n}{k_v N\tilde{F}\kappa B_n + \nu}, \\
A20' &= c_{prod} k_v N\tilde{F}\kappa B_n - c_{deg} A20, \qquad (12) \\
I\kappa B\alpha' &= c_{4a} I\kappa B\alpha_t - a_2 IKKa \cdot I\kappa B\alpha - \frac{a_3}{k_v} IKKa (C_T - k_v N\tilde{F}\kappa B_n) \frac{I\kappa B\alpha}{I\kappa B\alpha + \mu} \\
& \quad - c_{5a} I\kappa B\alpha - i_{1a} I\kappa B\alpha \frac{k_v N\tilde{F}\kappa B_n}{k_v N\tilde{F}\kappa B_n + \nu}, \\
I\kappa B\alpha_t' &= c_{1a} k_v N\tilde{F}\kappa B_n - c_{3a} I\kappa B\alpha_t.
\end{aligned}
$$

We re-write the system (12) as

$$
\begin{aligned}
IKKn' &= k_{prod} - (k_{deg} + T_R k_1) IKKn, \\
IKKa' &= T_R k_1 IKKn - k_3 IKKa - T_R k_2 IKKa \cdot A20 - k_{deg} IKKa, \\
NF\kappa B_n' &= a_3 IKKa \cdot (C_0 - NF\kappa B_n) \frac{\mu}{I\kappa B\alpha + \mu} - i_{1a} I\kappa B\alpha \frac{NF\kappa B_n}{NF\kappa B_n + \epsilon}, \\
A20' &= C_{prod} NF\kappa B_n - c_{deg} A20, \qquad (13) \\
I\kappa B\alpha' &= c_{4a} I\kappa B\alpha_t - a_2 IKKa \cdot I\kappa B\alpha - a_3 IKKa (C_0 - NF\kappa B_n) \frac{I\kappa B\alpha}{I\kappa B\alpha + \mu} \\
& \quad - c_{5a} I\kappa B\alpha - i_{1a} I\kappa B\alpha \frac{NF\kappa B_n}{NF\kappa B_n + \epsilon}, \\
I\kappa B\alpha_t' &= C_{1a} NF\kappa B_n - c_{3a} I\kappa B\alpha_t,
\end{aligned}
$$

where

$$C_0 := C_T/k_v,\ C_{prod} := c_{prod} k_v,\ C_{1a} := c_{1a} k_v,\ \epsilon = \nu/k_v$$

and ˜ has been dropped.

Next we introduce the following scaling

$$
\begin{aligned}
IKKn = \tfrac{k_{prod}}{k_{deg}} I\tilde{K}Kn, \quad IKKa = \tfrac{k_{prod}}{k_{deg}} I\tilde{K}Ka, \quad NF\kappa B_n = C_0 N\tilde{F}\kappa B_n, \\
A20 = \tfrac{C_{prod} C_0}{c_{deg}} \tilde{A20}, \qquad I\kappa B\alpha = C_0 I\kappa\tilde{B}\alpha, \qquad I\kappa B\alpha_t = \tfrac{C_{1a} C_0}{c_{3a}} I\kappa\tilde{B}\alpha_t,
\end{aligned} \qquad (14)
$$

where ˜ denotes the non-dimensional variable, and will be omitted from here on. Inserting

(14) into system (13) and dropping $\tilde{}$, we obtain

$$\begin{aligned}
IKKn' &= k_{deg}(1-IKKn)-T_R k_1 IKKn,\\
IKKa' &= T_R k_1 IKKn-(k_3+k_{deg}+T_R\bar{k}_2 A20)IKKa,\\
NF\kappa B_n' &= \bar{a}_3 IKKa\cdot(1-NF\kappa B_n)\frac{\delta}{I\kappa B\alpha+\delta}-i_{1a}I\kappa B\alpha\frac{NF\kappa B_n}{NF\kappa B_n+\varepsilon},\\
A20' &= c_{deg}(NF\kappa B_n-A20),\\
I\kappa B\alpha' &= \bar{c}_{4a}I\kappa B\alpha_t-\bar{a}_2 IKKa\cdot I\kappa B\alpha-\bar{a}_3 IKKa(1-NF\kappa B_n)\frac{I\kappa B\alpha}{I\kappa B\alpha+\delta}\\
&\quad -c_{5a}I\kappa B\alpha-i_{1a}I\kappa B\alpha\frac{NF\kappa B_n}{NF\kappa B_n+\varepsilon},\\
I\kappa B\alpha_t' &= c_{3a}(NF\kappa B_n-I\kappa B\alpha_t),
\end{aligned}\tag{15}$$

where

$$\begin{aligned}
&\bar{k}_2 := k_2\tfrac{c_{prod}}{c_{deg}}C_T, \qquad \delta := \tfrac{\mu}{C_0}, \qquad \varepsilon := \tfrac{\epsilon}{C_0}=\tfrac{\nu}{C_T},\\
&\bar{a}_2 := a_2\tfrac{k_{prod}}{k_{deg}}, \qquad \bar{a}_3 := a_3\tfrac{k_{prod}}{k_{deg}}, \qquad \bar{c}_{4a} := \tfrac{c_{4a}}{c_{3a}}C_{1a}.
\end{aligned}\tag{16}$$

In the consequence of nondimensionalization the number of parameters has been reduced from 18 to 13, see Table E. For convenience , we will drop $\bar{}$ in the notation of the parameters defined in (16), but it is to be understood that the parameters $k_2, a_2, a_3, c_{4a}$ of the reduced rescaled model are appropriately scaled versions of the corresponding parameters of the original model.

**Table E. Parameters of the rescaled reduced model (15) and their numerical values.**

| Parameter | Value | Unit | Description |
|---|---|---|---|
| $k_{deg}$ | 0.000125 | $s^{-1}$ | degradation of IKKn and IKKa |
| $k_1$ | 0.0025 | $s^{-1}$ | activation of IKKn caused by TNF |
| $k_3$ | 0.0015 | $s^{-1}$ | spontaneous inactivation of IKKa |
| $k_2$ | 0.0625 | $s^{-1}$ | inactivation of IKKa induced by A20 |
| $a_3$ | 0.2 | $s^{-1}$ | IKK (I$\kappa$B$\alpha$ \| NF-$\kappa$B) association leading to I$\kappa$B$\alpha$ degradation |
| $\delta$ | 0.0833 | | Michaelis type constant |
| $\varepsilon$ | 0.01667 | | Michaelis type constant |
| $c_{deg}$ | 0.000171 | $s^{-1}$ | effective A20 degradation |
| $c_{4a}$ | 0.0031 | $s^{-1}$ | I$\kappa$B$\alpha$ translation |
| $a_2$ | 0.04 | $s^{-1}$ | I$\kappa$B$\alpha$ degradation induced by IKKa |
| $c_{5a}$ | 0.0001 | $s^{-1}$ | I$\kappa$B$\alpha$ degradation |
| $i_{1a}$ | 0.001 | $s^{-1}$ | I$\kappa$B$\alpha$ nuclear import leading to NF-$\kappa$B removal from the nucleus |
| $c_{3a}$ | 0.0004 | $s^{-1}$ | I$\kappa$B$\alpha$ transcription and mRNA degradation |
