## Supplementary material for "A plausible identifiable model of the canonical NF-*κ*B signaling pathway": Suplementary Figures and Tables

featuring the article

### List of Tables

S1 **Summary of the considered stimulation protocols of the NF-$\kappa$B system in WT and A20 KO cells used for *in silico* experiments**. Each table block describes one TNF stimulation protocol for which the measured variables are: IKKa, nuclear NF-$\kappa$B, A20, total I$\kappa$B$\alpha$ protein denoted by I$\kappa$B$\alpha^*$ ($I\kappa B\alpha^* = I\kappa B\alpha + 1 - NF\kappa B$), and I$\kappa$B$\alpha$ mRNA. For each variable the measurement times used for the sensitivity-based identifiability analysis and reduced model fitting are given. . . . . 2

S2 **List of protocol sets, the corresponding numbers of measurement time points (N), and sensitivity vectors dimensions (dim).** Protocol sets are joint for WT and A20 KO. The combination experiment consists of 6 protocols: continuous and all pulsatile ones. The scaling factor is $dim^{0.5}$. . . . . 3

S3 **Parameter sets of the original model.** Five parameter sets of the original model randomly selected by varying the original parameters at most threefold from their original values. . . . . 4

S4 **Fitted parameter sets of the reduced model obtained from fitting to the original model with varied parameter sets**. . . . . 5

**Table S1. Summary of the considered stimulation protocols of the NF-$\kappa$B system in WT and A20 KO cells used for *in silico* experiments**. Each table block describes one TNF stimulation protocol for which the measured variables are: IKKa, nuclear NF-$\kappa$B, A20, total I$\kappa$B$\alpha$ protein denoted by I$\kappa$B$\alpha^*$ ($I\kappa B\alpha^* = I\kappa B\alpha + 1 - NF\kappa B$) , and I$\kappa$B$\alpha$ mRNA. For each variable the measurement times used for the sensitivity-based identifiability analysis and reduced model fitting are given.

| | |
|---|---|
| **continuous TNF *ON* 0-240min** | Reference: [1] |
| IKKa | 0, 5, 10, 15, 30, 60 min |
| NF-$\kappa$B, I$\kappa$B$\alpha^*$ | 0, 5, 10, 15, 30, 45, 60, 90, 120, 240 min |
| A20, I$\kappa$B$\alpha_t$ | 0, 15, 30, 45, 60, 90, 120, 240 min |
| **pulse 5-60 TNF *ON* 0-5 min, 60-65 min, 120-125 min** | Reference: [1] |
| IKKa | 0,5, 10, 15, 30, 60, 65, 70, 75, 90, 120, 125, 130, 135, 150, 180 min |
| NF-$\kappa$B, I$\kappa$B$\alpha^*$ | 0,5, 15, 30, 45, 60, 75, 90, 105, 120, 135, 150, 165, 180, 210, 240, 300, 360, 480, 720 min |
| A20, I$\kappa$B$\alpha_t$ | 0,5, 15, 30, 45, 60, 75, 90, 105, 120, 135, 150, 165, 180, 210, 240, 300, 360 min |
| **pulse 5-100 TNF *ON* 0-5 min, 100-105 min, 200-205 min** | Reference: [1] |
| IKKa | 0, 5, 10, 15, 30, 60, 105, 110, 115, 130, 160, 205, 210, 215, 230, 260 min |
| NF-$\kappa$B, I$\kappa$B$\alpha^*$ | 0, 5, 15, 30, 45, 60, 90, 115, 120, 130, 145, 160, 190, 215, 220, 230, 240, 245, 260, 290, 320, 340, 440, 560, 720 min |
| A20, I$\kappa$B$\alpha_t$ | 0, 5, 15, 30, 45, 60, 90, 115, 120, 130, 145, 160, 190, 215, 220, 230, 240, 245, 260, 290, 320, 340, 440 min |
| **pulse 5-200 TNF *ON* 0-5 min, 200-205 min, 400-405 min** | Reference: [1] |
| IKKa | 0, 5, 10, 15, 30, 60, 205, 210, 215, 230, 260, 405, 410, 415, 430, 460 min |
| NF-$\kappa$B, I$\kappa$B$\alpha^*$ | 0, 5, 15, 30, 45, 60, 90, 120, 215, 230, 240, 245, 260, 290, 320, 415, 430, 440, 445, 460, 490, 520, 640, 720 min |
| A20, I$\kappa$B$\alpha_t$ | 0, 5, 15, 30, 45, 60, 90, 120, 215, 230, 240, 245, 260, 290, 320, 415, 430, 440, 445, 460, 490, 520, 640 min |
| **pulse 22.5-45 TNF *ON* 0-22.5 min, 45-67.5 min, 90-112.5 min** | Reference: [2] |
| IKKa | 0, 5, 10, 15, 30, 50, 55, 60, 75, 95, 100, 105, 120, 150 min |
| NF-$\kappa$B, I$\kappa$B$\alpha^*$ | 0, 15 30, 45, 60, 75, 90, 105, 120, 135, 150, 165, 180, 210, 240, 285, 330, 450, 720 min |
| A20, I$\kappa$B$\alpha_t$ | 0, 15 30, 45, 60, 75, 90, 105, 120, 135, 150, 165, 180, 210, 240, 285, 330 min |
| **pulse 45-90 TNF *ON* 0-45 min, 90-135 min, 180-225 min** | Reference: [2] |
| IKKa | 0, 5, 10, 15, 30, 60, 95, 100, 105, 120, 150, 185, 190, 195, 210, 240 min |
| NF-$\kappa$B, I$\kappa$B$\alpha^*$ | 0, 15, 30, 45, 60, 90, 105, 120, 135, 150, 180, 195, 210, 225, 240, 270, 300, 330, 420, 540, 720 min |
| A20, I$\kappa$B$\alpha_t$ | 0, 15, 30, 45, 60, 90, 105, 120, 135, 150, 180, 195, 210, 225, 240, 270, 300, 330, 420 min |

**Table S2. List of protocol sets, the corresponding numbers of measurement time points (N), and sensitivity vectors dimensions (dim).** Protocol sets are joint for WT and A20 KO. The combination experiment consists of 6 protocols: continuous and all pulsatile ones. The scaling factor is $dim^{0.5}$.

| **Protocol sets** | **N** | **dim** | **dim$^{0.5}$** |
|---|---|---|---|
| continuous | 76 | 67 | 8.19 |
| pulse 5-60 | 166 | 157 | 12.53 |
| pulse 5-100 | 201 | 192 | 13.86 |
| pulse 5-200 | 197 | 188 | 13.71 |
| pulse 22.5-45 | 155 | 146 | 12.08 |
| pulse 45-90 | 173 | 164 | 12.81 |
| continuous<br>pulse 5-60 | 242 | 224 | 14.97 |
| continuous<br>pulse 5-100 | 277 | 259 | 16.09 |
| continuous<br>pulse 5-200 | 273 | 255 | 15.97 |
| continuous<br>pulse 22.5-45 | 231 | 213 | 14.59 |
| continuous<br>pulse 45-90 | 249 | 231 | 15.20 |
| combination experiment | 968 | 914 | 30.23 |
| on-off | 50 | 41 | 6.40 |

**Table S3. Parameter sets of the original model.** Five parameter sets of the original model randomly selected by varying the original parameters at most three-fold from their original values.

| Parameter | Set 1 | Set 2 | Set 3 | Set 4 | Set 5 |
|---|---|---|---|---|---|
| $c_1$ | 6.74E-07 | 7.04E-07 | 1.85E-07 | 3.31E-07 | 5.61E-07 |
| $c_3$ | 9.13E-04 | 1.96E-04 | 7.11E-04 | 1.92E-04 | 3.04E-04 |
| $c_4$ | 4.71E-01 | 2.45E-01 | 8.66E-01 | 1.33E+00 | 3.06E-01 |
| $c_5$ | 4.88E-04 | 1.30E-04 | 8.78E-04 | 5.73E-04 | 2.75E-04 |
| $k_1$ | 2.04E-03 | 5.85E-03 | 2.11E-03 | 8.48E-04 | 4.65E-03 |
| $k_2$ | 1.01E-01 | 3.90E-02 | 8.93E-02 | 2.57E-01 | 2.94E-01 |
| $k_3$ | 5.60E-04 | 8.46E-04 | 9.50E-04 | 7.88E-04 | 7.92E-04 |
| $k_{prod}$ | 1.50E-05 | 3.57E-05 | 1.70E-05 | 1.63E-05 | 2.16E-05 |
| $k_{deg}$ | 1.10E-04 | 6.74E-05 | 8.61E-05 | 1.44E-04 | 7.48E-05 |
| $a_2$ | 1.87E-01 | 2.82E-01 | 1.26E-01 | 1.11E-01 | 1.32E-01 |
| $a_1$ | 8.33E-01 | 4.67E-01 | 5.35E-01 | 9.84E-01 | 9.70E-01 |
| $a_3$ | 8.00E-01 | 2.54E+00 | 1.33E+00 | 1.33E+00 | 3.34E-01 |
| $t_1$ | 6.83E-02 | 2.02E-01 | 4.43E-02 | 1.89E-01 | 2.14E-01 |
| $t_2$ | 2.42E-01 | 4.88E-02 | 1.36E-01 | 2.12E-01 | 1.33E-01 |
| $c_{1a}$ | 1.01E-06 | 2.76E-07 | 3.49E-07 | 5.39E-07 | 1.13E-06 |
| $c_{3a}$ | 8.63E-04 | 4.91E-04 | 2.07E-04 | 9.36E-04 | 4.29E-04 |
| $c_{4a}$ | 3.83E-01 | 5.47E-01 | 1.13E+00 | 9.67E-01 | 1.42E+00 |
| $c_{5a}$ | 2.69E-04 | 2.91E-04 | 1.12E-04 | 1.06E-04 | 1.05E-04 |
| $c_{6a}$ | 1.33E-05 | 1.11E-05 | 1.14E-05 | 3.74E-05 | 7.45E-06 |
| $i_1$ | 4.65E-03 | 6.45E-03 | 2.57E-03 | 9.69E-04 | 5.32E-03 |
| $e_{2a}$ | 3.48E-03 | 3.67E-03 | 4.53E-03 | 1.03E-02 | 9.59E-03 |
| $i_{1a}$ | 3.76E-04 | 4.67E-04 | 6.66E-04 | 7.86E-04 | 1.24E-03 |
| $e_{1a}$ | 5.62E-04 | 1.07E-03 | 2.16E-04 | 4.78E-04 | 4.68E-04 |

**Table S4. Fitted parameter sets of the reduced model obtained from fitting to the original model with varied parameter sets**.

| Parameter | Fit 1 | Fit 2 | Fit 3 | Fit 4 | Fit 5 |
|---|---|---|---|---|---|
| $a_2$ | 2.43E-02 | 1.19E-01 | 4.69E-02 | 2.62E-02 | 1.65E-02 |
| $a_3$ | 7.10E-02 | 4.04E-01 | 1.71E-01 | 5.03E-02 | 9.17E-02 |
| $c_{4a}$ | 2.57E-03 | 2.18E-03 | 7.94E-03 | 2.78E-03 | 1.35E-02 |
| $\delta$ | 7.92E-02 | 5.32E-01 | 3.92E-02 | 3.86E-02 | 8.55E-03 |
| $\varepsilon$ | 2.96E-02 | 1.67E-01 | 7.51E-02 | 1.67E-03 | 2.83E-02 |
| $k_2$ | 1.59E-02 | 1.28E-01 | 6.25E-03 | 8.55E-02 | 1.29E-01 |
| $c_{3a}$ | 7.44E-04 | 4.00E-04 | 1.94E-04 | 7.61E-04 | 3.50E-04 |
| $c_{5a}$ | 4.24E-04 | 1.00E-03 | 4.04E-05 | 4.19E-05 | 6.04E-05 |
| $c_{deg}$ | 2.56E-04 | 1.71E-05 | 3.02E-04 | 6.89E-05 | 6.82E-05 |
| $i_{1a}$ | 3.62E-04 | 7.30E-04 | 4.59E-04 | 3.56E-04 | 6.82E-04 |
| $k_1$ | 1.84E-03 | 9.44E-03 | 2.12E-03 | 9.37E-04 | 4.97E-03 |
| $k_3$ | 4.72E-04 | 7.25E-04 | 9.30E-04 | 7.20E-04 | 8.11E-04 |
| $k_{deg}$ | 8.04E-05 | 8.33E-05 | 9.02E-05 | 1.20E-04 | 4.81E-05 |

S4 **Practical identifiability of the reduced model based on Monte Carlo simulations.** A comparison of 75% confidence ellipses (shown in black) computed for $\sigma_{data} = 1.3$ in the linear sensitivity matrix based analysis with results from 50 Monte Carlo simulations for four values of $\sigma_{data}$ (1.0, 1.1, 1.2, 1.3). Shown are projections on 21 planes spanned by 7 parameters with the smallest geometric standard deviations $\sigma_{carlo,j}$ (for $\sigma_{data} = 1.3$). . . . . . . . . 10

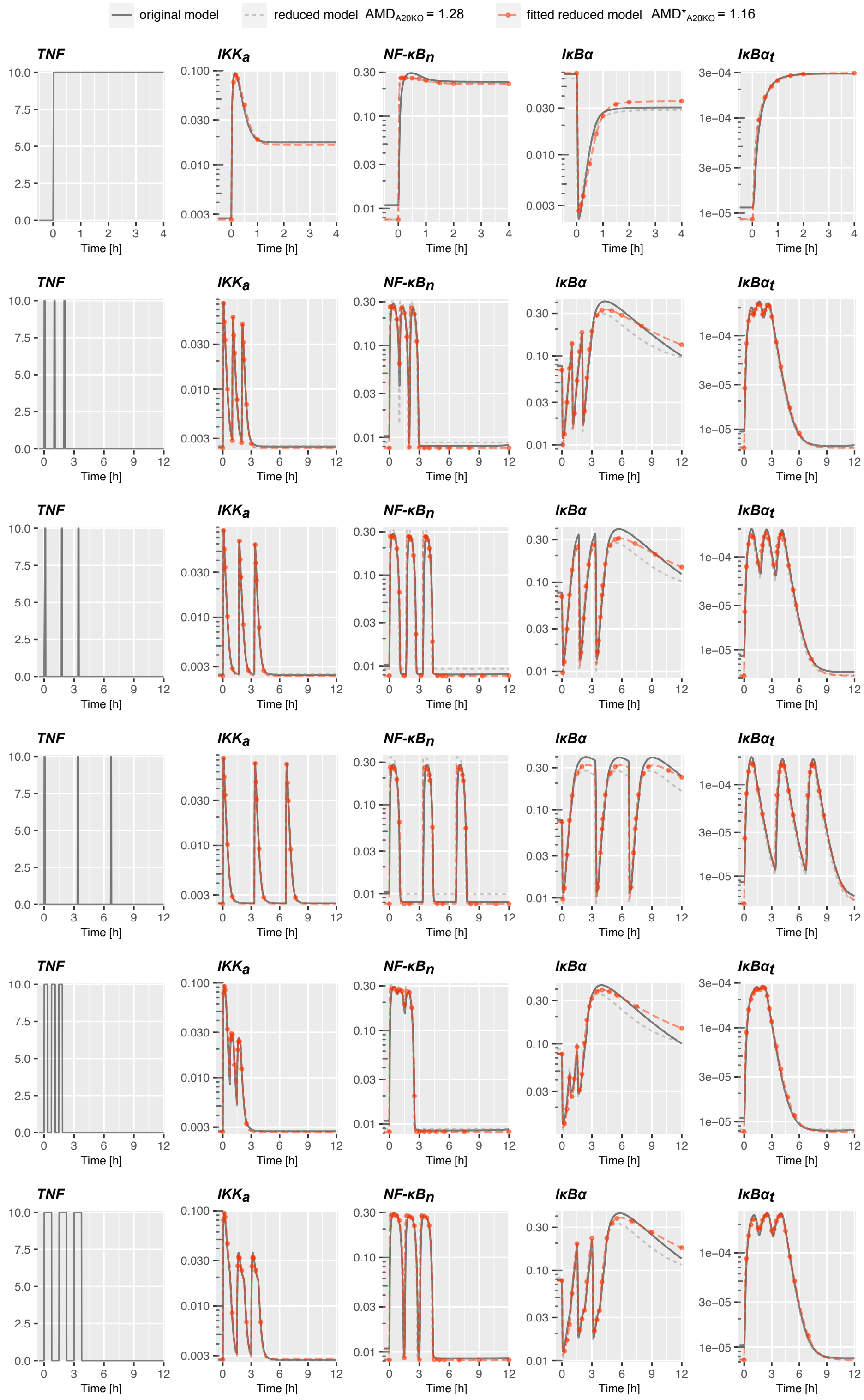

**Fig S1. Dynamics of the original, reduced and reduced fitted models in combination experiment in A20 KO cells defined in Table S1.** Red dots indicate time points from which the nuclear NF-κB and total IκB$\alpha$ protein *in silico* measurements are used for fitting the reduced model to the original one. The time points for the remaining variables are given in Table S1.

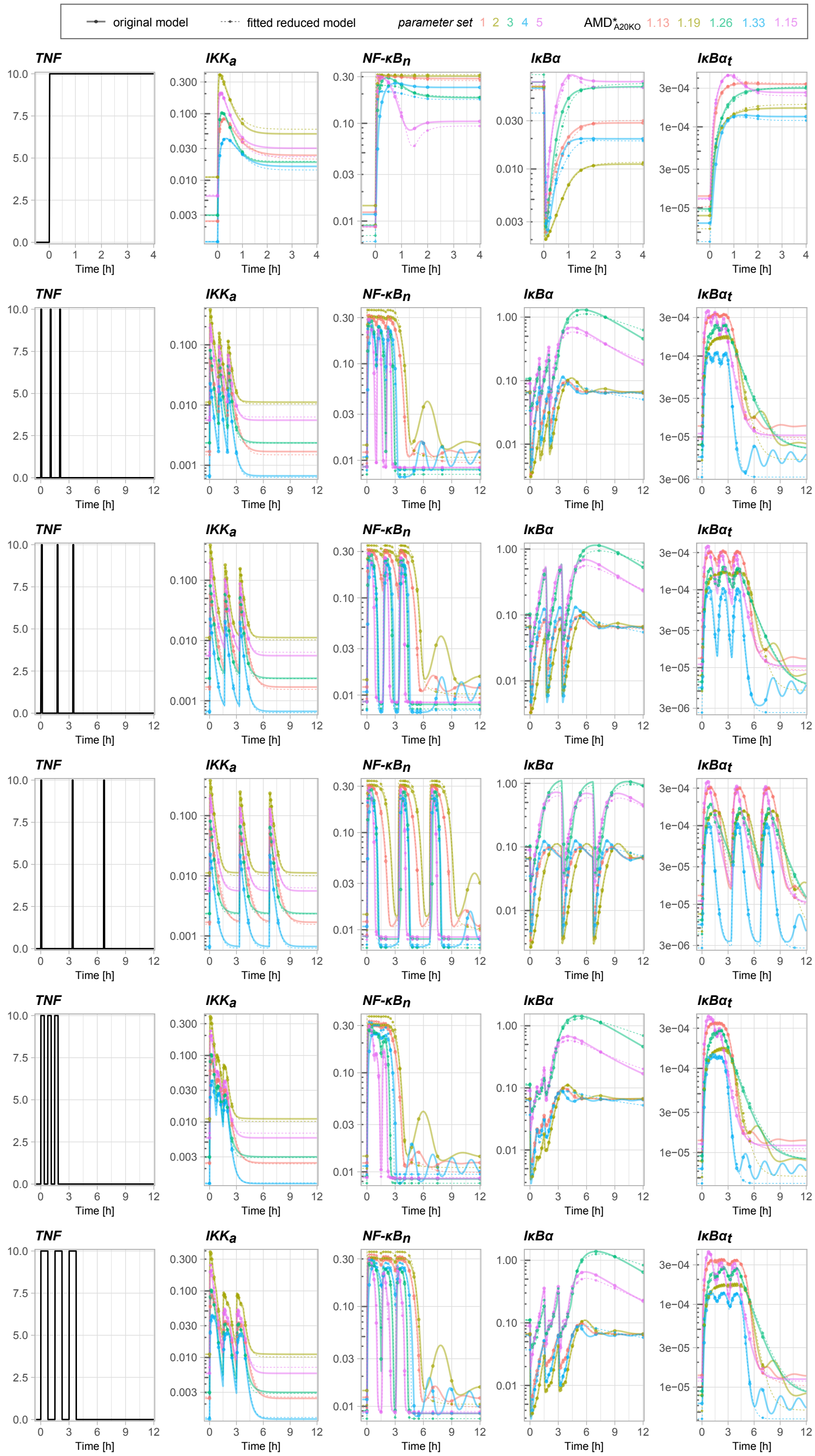

**Fig S2. Simulations of the original model for A20 KO cells for five different sets of parameters and the corresponding reduced model with refitted parameter values (see Table S3 and Table S4** Simulations are performed for the combination experiment defined in Table S1. For each parameter set the corresponding $AMD^*_{A20KO}$ is computed based on trajectories of the 5 main model variables.

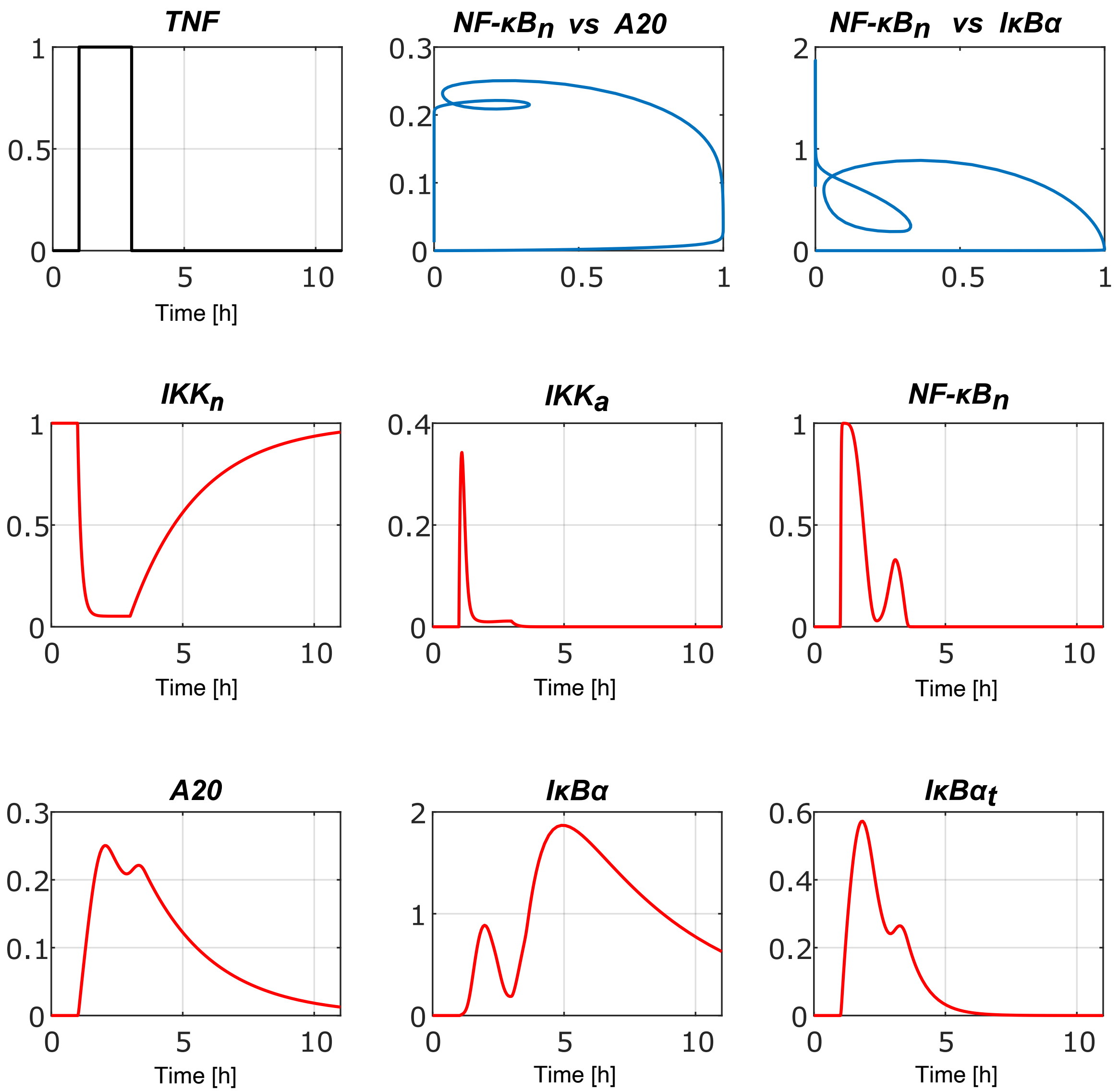

**Fig S3. Dynamics of the reduced fitted model for the on-off protocol in WT cells defined in Table 2.**

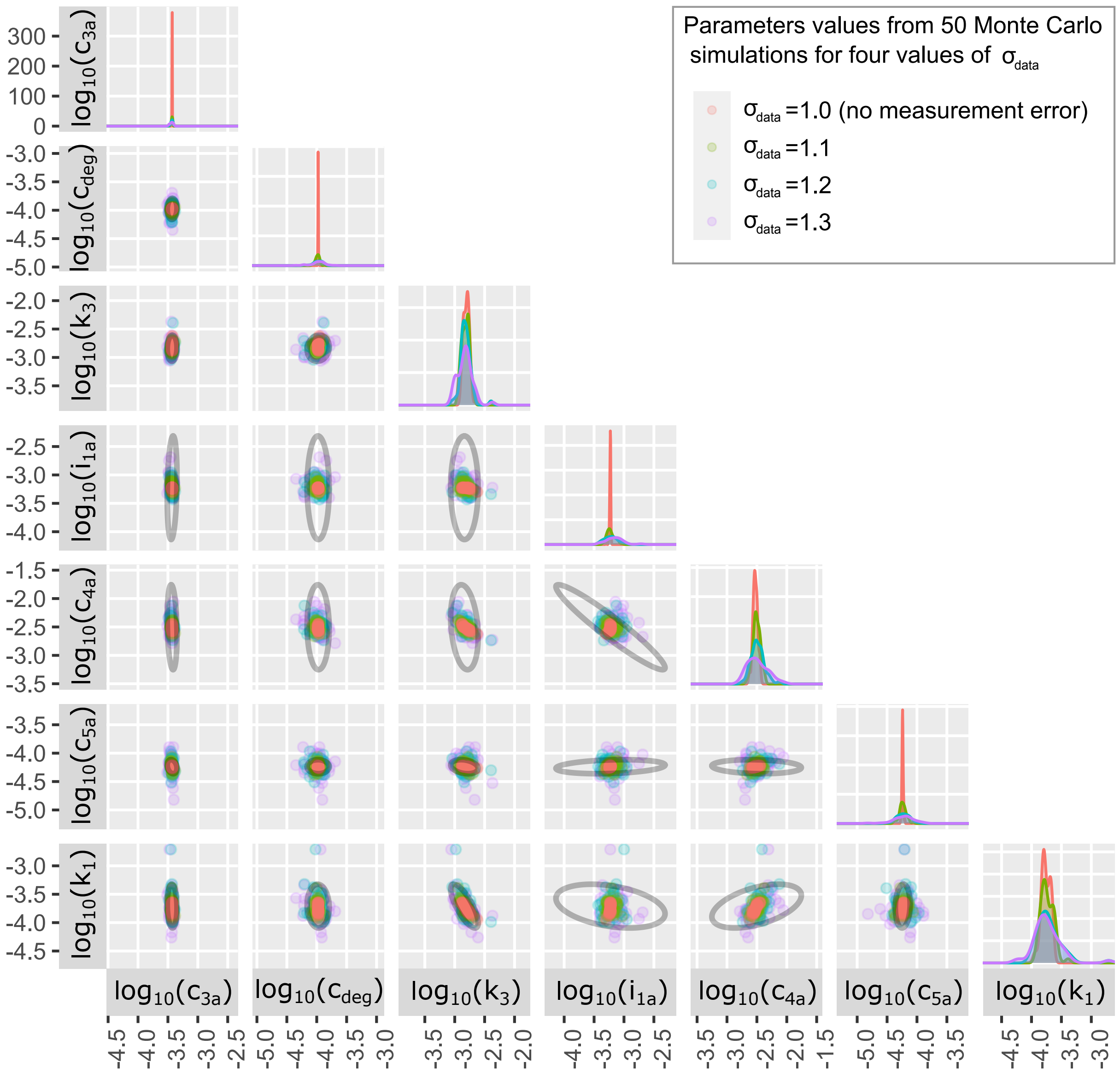

**Fig S4. Practical identifiability of the reduced model based on Monte Carlo simulations.** A comparison of 75% confidence ellipses (shown in black) computed for $\sigma_{data} = 1.3$ in the linear sensitivity matrix based analysis with results from 50 Monte Carlo simulations for four values of $\sigma_{data}$ (1.0, 1.1, 1.2, 1.3). Shown are projections on 21 planes spanned by 7 parameters with the smallest geometric standard deviations $\sigma_{carlo,j}$ (for $\sigma_{data} = 1.3$).

[2] Zambrano S, De Toma I, Piffer A, Bianchi ME, Agresti A. NF-$\kappa$B oscillations translate into functionally related patterns of gene expression. eLife 5:e09100. doi:10.7554/eLife.09100
