## Supplementary Codes - zipped directory for "A plausible identifiable model of the canonical NF-*κ*B signaling pathway": Jaruszewicz_et_al_BNGL_README.txt

This directory contains BNGL files with the original and reduced NF-kB models developed in the paper

"A plausible identifiable model of the canonical NF-κB signaling pathway" 

by Joanna Jaruszewicz-Blonska, Ilona Kosiuk, Wiktor Prus, and Tomasz Lipniacki


======================================================

File list:

nfkb_fitted_reduced_model_combination_experiment.bngl

nfkb_fitted_reduced_model_on_off.bngl

NFkB_original_model_combination_experiment.bngl

======================================================

Description:


Each BNGL file describes the model specification and a set of commands that operate on the model.


- - - 1 - - - nfkb_fitted_reduced_model_combination_experiment.bngl

This file describes the reduced NF-kB model.
The parameter values are taken from Table 1 (fitted values). 
The simulation shows the dynamics of the (fitted) reduced model in the combination experiment (defined in S1 Table)  for WT cells (shown in red in Fig 2) and A20KO cells (S1 Fig). 

The parameters can be changed to pre-fitted values (Table 1) to reproduced the dynamics of the rescaled reduced model (before its fitting to the original model), shown in Fig 2 (grey dashed). 
Changing the parameters to one of 5 parameter sets from S4 Table, reproduces the results shown in Fig 3 for WT cells and results in S2 Fig for A20KO cells.

  

- - - 2 - - - nfkb_fitted_reduced_model_on_off.bngl

This file describes the reduced NF-kB model and the on-off protocol (defined in Table 2).
The parameter values are taken from Table 1 (fitted values). 
The simulation shows the dynamics of the (fitted) reduced model for the on-off protocol (defined in Table 2)  for WT cells (shown in S3 Fig) and A20KO cells. 


- - - 3 - - - NFkB_original_model_combination_experiment.bngl

This file describes the original NF-kB model.
The parameter values are taken from Table A in S1_Supplementary_Text.  The simulation shows the dynamics of the original model in the combination experiment (defined in S1 Table) for WT cells (shown in black in Fig 2) and A20KO cells (S1 Fig). 

Changing the parameters to one of 5 parameter sets from S3 Table, reproduces the results shown in Fig 3 for WT cells and results in S2 Fig for A20KO cells.


============================


How to:

 0.    Unzip the file “BNGL_files.zip". (A directory “BNGL_files" is generated).
 1.    Start RuleBender.
 2.    Run one of the files: 

       nfkb_fitted_reduced_model_combination_experiment.bngl
       nfkb_fitted_reduced_model_on_off.bngl
       NFkB_original_model_combination_experiment.bngl

 3.    The new parameters can be specified in each of the bngl files.

 

# For a description of the BioNetGen language (BNGL) see e.g. Faeder et al
# [Meth. Mol. Biol. (2009), http://dx.doi.org/10.1007/978-1-59745-525-1_5].
# The model may be executed using BioNetGen [http://www.bionetgen.org].
# We recommend using BioNetGen within RuleBender [http://www.rulebender.org].


 
