## Supplementary Codes - zipped directory for "A plausible identifiable model of the canonical NF-*κ*B signaling pathway": Jaruszewicz_et_al_README.txt

This directory contains MATLAB files with the reduced NF-kB model developed in the paper

"A plausible identifiable model of the canonical NF-κB signaling pathway" 

by Joanna Jaruszewicz-Blonska, Ilona Kosiuk, Wiktor Prus, and Tomasz Lipniacki


====================

File list:

model.m
program_continuous.m
program_5min_60min.m
program_on_off.m

=====================

Description:

- - - 1 - - - model.m

This MATLAB function characterizes a system of ODEs describing the reduced NF-kB model.
The parameter values are taken from Table 1 (fitted values) but can be changed in order to obtain different types of behaviour (e.g. as shown in Fig 8 and Fig 9). 


- - - 2 - - - program_continuous.m

This is a master program (calling model.m) for the reduced NF-kB model for the protocol “continuous” described in S1 Table.
Running the program one can reproduce the results presented in Fig 2 (first row), i.e., numerical trajectories of wild type cells of the 5 main variables (IKKa, NF-κB, A20, total IκBα, and mRNA IκBα), generated by the reduced fitted model (shown in red). 


- - - 3 - - - program_5min_60min.m

This is a master program (calling model.m) for the reduced NF-kB model for the protocol “pulse 5-60” described in S1 Table.
Running the program one can reproduce the results presented in Fig 2 (second row), i.e., numerical trajectories of wild type (WT) cells of the 5 main variables (IKKa, NF-κB, A20, total IκBα, and mRNA IκBα), generated by the reduced fitted model (shown in red). 


- - - 4 - - - program_on_off.m

This is a master program (calling model.m) for the reduced NF-kB model for the protocol “on-off” described in Table 2.
Running the program one can reproduce the results presented in S3 Fig, i.e., numerical trajectories of wild type (WT) cells of the 5 main variables (IKKa, NF-κB, A20, total IκBα, and mRNA IκBα), generated by the reduced fitted model (shown in red). 

============================


How to:

 0.    Unzip the file “MATLAB_codes.zip". (A directory “MATLAB_codes" is generated).
 1.    Start MATLAB.
 2.    Run one of the files: program_continuous.m,  program_5min_60min.m  or  program_on_off.m  
 3.    The new parameters can be specified in the ‘model.m’ file.

 


 
